# The adeno-associated virus Rep proteins target PP4:SMEK1 by preventing substrate-recruitment

**DOI:** 10.1101/2025.06.13.659448

**Authors:** Bram Vandewinkel, Sophie Torrekens, Zander Claes, Mathieu Bollen, Els Henckaerts

## Abstract

Adeno-associated virus (AAV) vectors are widely used for in vivo gene therapy, but their clinical effectiveness and scalable production remain limited by gaps in our understanding of AAV biology. In particular, the host–virus interactions that regulate AAV gene expression and replication are not fully defined, yet they hold key insights for engineering improved recombinant AAV (rAAV) vectors. Here, we identify the PP4:SMEK1/2 phosphatase complex as a host restriction factor that suppresses wildtype AAV replication. Active replication and Rep expression correlates with hyperphosphorylation of KAP1^S824^ and RPA2^S4/8/33^, known PP4 substrates. Rep proteins directly bind the HEAT/Arm domain of the SMEK1 regulatory subunit of PP4. We further reveal that the KAP1:SMEK1 interaction is mediated by a MAPP short linear motif (SLiM) that interacts with the substrate recruitment groove of the SMEK1 EVH1 domain. Rep68 disrupts this interaction and interferes with substrate recruitment by the PP4:SMEK1/2 complex, leading to substrate hyperphosphorylation. We also uncover a multifunctional complex containing PP4:SMEK1 and PP1:NIPP1, both implicated in KAP1^S824^ dephosphorylation. Together, these findings reveal a substrate-recruitment interference mechanism by which AAV Rep proteins inhibit host phosphatase activity. This mechanistic insight provides a blueprint for enhancing rAAV vector potency and production in gene therapy applications.

## Introduction

Protein phosphatases are essential regulators of cellular signaling, serving to balance the activity of protein kinases. Once considered mere housekeeping components, they are now recognized as integral regulators of diverse cellular pathways and have been implicated in the pathogenesis of numerous diseases ^1^. Given their role in cellular processes such as cell cycle regulation, transcription, protein synthesis and apoptosis, it is not surprising that viruses have evolved mechanisms to hijack cellular phosphatases to benefit their replication. A growing body of research highlights the critical role of reversible protein phosphorylation in the replication of viruses, and both phosphorylation and dephosphorylation of viral and cellular proteins have been shown to play key roles in supporting the life cycle of many viruses. Viral interactions with serine/threonine phosphatase PP2A for example serve to inhibit PP2A-mediated dephosphorylation of various cellular substrates, leading to cell cycle arrest (HIV-Vif) or transformation of the infected cell ^2–8^. The viral mechanism of PP2A inhibition ranges from degradation of PP2A’s regulatory subunit B56 ^9–11^, as described for HIV, to PP2A complex binding and remodeling as seen in SV40, HIV and HBV replication. Conversely, some viruses upregulate or stimulate PP2A activity to suppress innate immune signaling ^12–14^. EBOV on the other hand exploits PP2A to dephosphorylate the viral protein VP30 and kickstart viral transcription ^15^. Similarly, viruses exploit serine/threonine protein phosphatase 1 (PP1) to promote viral transcription or to overcome the block in translation ^16,17^, mediated through viral RNA signaling or ER stress induced by viral protein expression. Additionally, a myriad of virus-host interactions involving PP1 have been described to control the innate immune response elicited by viral infection ^18^. Moreover, our previous studies on the interactions between the adeno-associated virus (AAV) and PP1 revealed that viruses also manipulate cellular phosphatases to counteract host cell-mediated chromatinization and silencing of the viral genome ^19^.

AAV is a non-pathogenic parvovirus with a 4.7 kb ssDNA viral genome, containing two major open reading frames (ORFs) ^20,21^. In total, nine different viral proteins are expressed during active replication ^21–23^, a process that requires the presence of a helper virus such as adenovirus or herpes simplex virus ^24,25^. Establishment of the lytic phase of the viral life cycle depends on the expression of various helper virus proteins and the AAV non-structural replication (Rep) proteins. These Rep proteins play an essential role in viral transcription, replication and packaging ^26–29^ and are expressed from the same ORF in the AAV genome through a complex interplay of alternative promoter usage and splicing events. The four AAV Rep isoforms—Rep78, Rep68, Rep52, and Rep40—share an AAA^+^-ATPase/helicase domain. Rep78 and Rep68 also have a N-terminal endonuclease domain, but only Rep78 and Rep52 have a C-terminal zinc finger. To extend their function beyond what is provided by their individual domains, these proteins assemble into various oligomeric states ^30,31^ and interact with host cellular proteins to optimally exploit and control the host cell ^32^.

Various protein-protein interaction approaches have been employed to reveal cellular interaction partners of the Rep proteins, however the molecular mechanisms and purpose of these interactions remain largely unexplored. Using an unbiased BioID approach to determine the putative Rep interactome, we discovered that the viral replication proteins interact with the PP1:NIPP1 holoenzyme and inhibit its activity ^19^. This inhibition maintains the transcriptional co-repressor KAP1^S824^ in a phosphorylated, inactive state, causing the dissociation of histone-modifying proteins (complexes) from KAP1, thereby facilitating viral transcription. Notably, dephosphorylation of KAP1^S824^ has also been reported to be regulated by protein phosphatase 4 (PP4) ^33^, a member of the same phosphatase family as PP1. Interestingly, the PP4 regulatory subunit SMEK1 (PP4R3A) was also identified as a putative Rep interactor ^19^.

PP4 is a serine/threonine-specific phosphatase belonging to the phosphoprotein phosphatase (PPP) family and is considered a PP2A-like phosphatase. Although the regulation of its activity, substrate specificity, and cellular localization appear less complex than those of PP1 and PP2A, PP4 remains relatively understudied. The PP4 catalytic subunit forms both heterodimeric complexes (with PP4R1 or PP4R4) and heterotrimeric complexes (with PP4R2 and either PP4R3A or PP4R3B). Among these, the heterotrimeric complexes are currently the best characterized. The regulatory subunits PP4R3A and PP4R3B, also known as SMEK1 and SMEK2, respectively, recruit substrates via their N-terminal EVH1 domains for dephosphorylation by associated PP4. These substrates typically contain a short linear motif (SLiM), either FxxP or MxPP, which binds to the EVH1 domain with low micromolar affinity ^34^. Despite this defined recruitment mechanism, only a limited number of validated PP4:SMEK1/2 substrates have been identified, including KAP1, RPA2 and γH2AX, suggesting that PP4 primarily functions in processes such as the DNA-damage response (DDR) and cell cycle regulation ^33,35,36^. Furthermore, the role of PP4 holoenzymes in the viral life cycle is largely unexplored, and to date, no viral interactions with PP4:SMEK complexes have been reported.

Here, we report for the first time that the PP4 phosphatase, in complex with its regulatory subunit SMEK1, is targeted by viral proteins. We demonstrate that PP4, together with SMEK1 and SMEK2, plays a repressive role in the AAV life cycle. To overcome this restriction, the viral Rep proteins interfere with PP4:SMEK activity, maintaining substrates such as KAP1 and RPA2 in a hyperphosphorylated state during lytic replication. We further show that AAV Rep proteins employ a novel, previously undescribed mechanism of PP4 substrate-recruitment interference. Finally, we identify a large multifunctional phosphatase complex composed of the PP1:NIPP1 and PP4:SMEK1 holoenzymes, suggesting that AAV Rep proteins simultaneously target multiple phosphatase pathways.

## Results

### AAV Rep proteins interact directly with SMEK1

To test our hypothesis that the AAV Rep proteins interact with the PP4:SMEK1 holoenzyme complex, we performed GFP-trap experiments with GFP-tagged SMEK1 from cells that were co-infected with AAV2 (10 IU) and Ad5 (MOI 5) (Figure 1A). Pull-down of GFP-SMEK1 at 28h post-infection (pi) without formaldehyde crosslinking resulted in robust co-precipitation of the PP4 catalytic subunit, all four Rep isoforms, as well as the PP4 substrate KAP1 ^33^, and SF3B1, a PP1:NIPP1 substrate not previously reported to associate with the PP4 holoenzyme (Figure 1B) ^37^. Formaldehyde crosslinking was employed because prior attempts to detect the interaction between Rep and the PP1:NIPP1 holoenzyme had proven challenging ^19^. The enzyme-substrate interaction between SMEK1 and KAP1 was further enhanced by prior crosslinking, whereas the SMEK1:Rep interaction diminished upon crosslinking, possibly caused by stabilization of more dynamic SMEK1 protein complexes (Figure 1B). This may indicate that Rep interacts stronger, and/or less dynamic with SMEK1 than with KAP1. Reciprocal pull-down assays with overexpression of the four different Rep isoforms confirmed the SMEK1:Rep interaction (Figure S1A).

**Figure 1.**
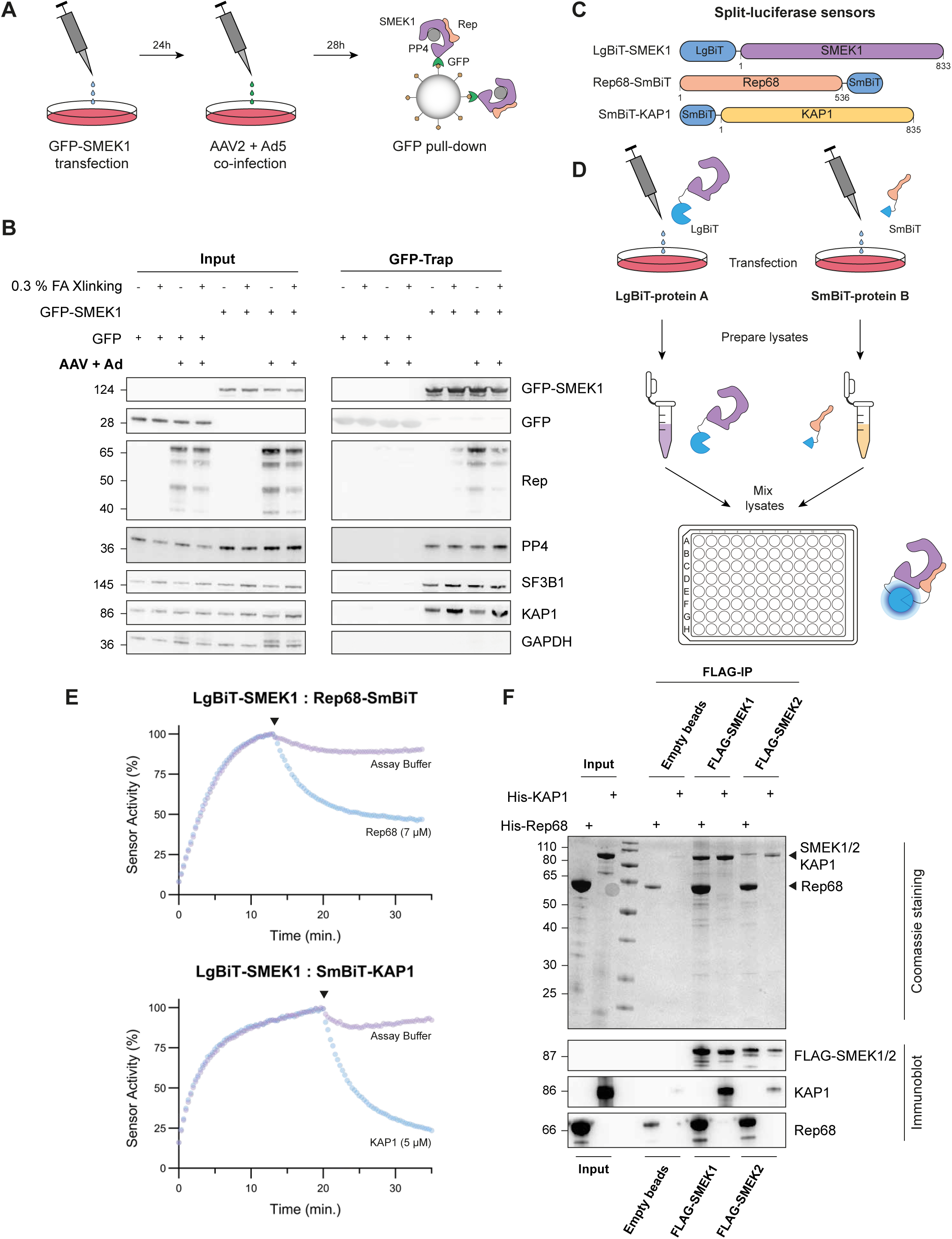
PP4R3A (SMEK1) directly interacts with the AAV Rep proteins. **(A)** Schematic representation of the GFP-trap experiment of GFP-tagged SMEK1 during a lytic AAV infection. **(B)** Co-immunoprecipitation of endogenous expressed Rep, PP4, SF3B1 and KAP1 with ectopically expressed GFP-SMEK1, in the presence/absence of formaldehyde crosslinking. GFP alone is included as a control to account for non-specific interactions. Input samples are shown on the left, while pull-down/trap samples are shown on the right. **(C)** Design of the split-luciferase sensors used for protein-protein interaction (PPI) studies. SMEK1 was N-terminally tagged with the LgBiT tag, while Rep68 and KAP1 were respectively C- or N-terminally tagged with the SmBiT tag. A flexible linker of 15-20 AA was included in between the tag and protein. **(D)** Experimental design of the lysate-based split-luciferase assays performed with the LgBiT-SMEK1:Rep68-SmBiT and LgBiT-SMEK1:SmBiT-KAP1 sensors as described before ^41^. Assays were performed in a white 384-well low protein binding plate. **(E)** Kinetic-trace experiment showing the time-dependent association of the LgBiT-SMEK1:Rep68-SmBiT and LgBiT-SMEK1:SmBiT-KAP1 split-luciferase sensors. The black arrow indicates the addition of the purified, untagged competitor (concentration indicated in the graph). The represented data is plotted as a percentage of the signal-to-background (S/B) ratio right before the addition of the competitor. The data shown is a representative example of three independent repeats. **(F)** FLAG-IP of purified FLAG-SMEK1 and FLAG-SMEK2 transiently expressed in HEK293T cells. Purified FLAG-SMEK1/2 bound to anti-FLAG agarose beads was incubated with purified His-tagged KAP1 or Rep68 to investigate co-precipitation. Co-immunoprecipitation of Rep68 and KAP1 was confirmed via Coomassie staining (top panel) and immunoblotting (bottom panel). Empty beads (beads incubated with non-transfected HEK293T lysate) served as a control to assess the stickiness of the purified Rep68 and KAP1.

To get more insight into the interaction dynamics, we next performed lysate-based split-luciferase assays. This assay, employed to study protein-protein interactions, relies on the separate fusion of two catalytically inactive NanoLuc fragments (LgBiT and SmBiT) to two proteins of interest. The lysate-based variant was recently developed and successfully used to study the interaction of various PP1 and PP2A holoenzymes ^38–41^. We fused SMEK1 to the LgBiT tag, while Rep68 and KAP1 were tagged with SmBiT, with KAP1 serving as a positive control (Figure 1C). Rep68 was selected as a representative for studying interactions between the Rep proteins and SMEK1, due to its ease of purification from *E. coli* with high yield. To study the dynamic behaviour of the SMEK1:Rep68 and SMEK1:KAP1 interaction, we prepared HEK293T cell lysates from cells that separately expressed LgBiT-SMEK1, Rep68-SmBiT and SmBiT-KAP1 (Figure 1D). To monitor the association of each interaction sensor, lysates containing LgBiT-SMEK1 were combined with either Rep68-SmBiT or SmBiT-KAP1, and luminescence was measured over time. To assess the specificity and dynamic behaviour of these interactions, we introduced purified His-tagged partner proteins lacking the SmBiT tag (i.e., His-Rep68 or His-KAP1, hereafter referred to as “untagged competitor”) as competitors (Figure S1B,C). The untagged competitors were added at relatively high concentrations (low micromolar) to effectively outcompete the SmBiT-tagged variants, leading to sensor dissociation (Figure 1D,E). Upon addition of untagged competitor, dissociation was observed for both interaction sensors (Figure 1E), strongly indicating that the interactions detected in the lysate-based split-luciferase assays are specific and dynamic.

Lastly, we extended the lysate-based assays with an *in vitro* pull-down approach to study the direct interaction of purified Rep68/KAP1 and SMEK1. Ectopically expressed FLAG-SMEK1 was purified from HEK293T cells. Next, the purified FLAG-SMEK1 bound to the beads was incubated with either recombinant Rep68 or KAP1. Following multiple washing steps, Rep68 and KAP1 co-precipitated with purified FLAG-SMEK1, suggesting a direct interaction with SMEK1 (Figure 1F). We also included FLAG-SMEK2-bound beads and observed that Rep68 also interacts strongly with the alternative PP4 regulatory subunit that constitutes the other heterotrimeric PP4 complex (Figure 1F). To further validate the direct interaction with SMEK1, we performed split-luciferase assays using recombinantly expressed and purified LgBiT-SMEK1, Rep68-SmBiT, and SmBiT-KAP1 (Figure S1B-D). Consistent with previous results, the purified interaction sensors showed clear association, which was reversed by addition of the untagged competitor (Figure S1D).

In summary, we demonstrated an interaction between the AAV Rep proteins and the PP4 regulatory subunit SMEK1. In addition, *in vitro* assays with purified proteins revealed that SMEK1 interacts directly with Rep68 and KAP1.

### Rep68 interacts with the HEAT/Arm domain of SMEK1, independent of PP4

To gain more insights in how PP4 and the Rep proteins bind to SMEK1, we made deletion mutants of SMEK1 (Figure 2A and S2A). The N-terminal EVH1 domain of SMEK1 plays a critical role in substrate recruitment by recognizing short linear motifs (SLiMs), specifically those containing FxxP or MxPP sequences ^34^. The HEAT/Arm domain functions as a scaffold for coordinating PP4 assembly ^42^, while its unstructured C-terminal region contains its nuclear localization signal (NLS) ^43^. SMEK1 truncations were tested for loss of PP4 binding using lysate-based split-luciferase assays. As expected, deletion of the HEAT/Arm domain of SMEK1 resulted in a complete loss of binding with PP4, whereas removal of the EVH1 domain did not affect this interaction (Figure 2B and C). These results were confirmed by GFP-SMEK1 pull-downs (Figure 2D). Surprisingly, removal of the unstructured C-terminus enhanced the interaction of PP4 with the HEAT/Arm domain, indicating it somehow hinders the PP4 coordination by SMEK1. We also examined the purified SMEK1 deletion mutants for binding to purified Rep68-SmBit, using split-luciferase assays. Binding of Rep68 to SMEK1 was decreased after deletion of either the EVH1 or HEAT/Arm domain (Figure S2A-C). However, stronger loss of binding was observed in the SMEK1 deletion mutants where the HEAT/Arm domain was removed, indicating that Rep68 mainly interacts with the latter domain. Pull-down assays further confirmed that the SMEK1:Rep68 interaction is mainly mediated by the HEAT/Arm domain (Figure 2D).

**Figure 2.**
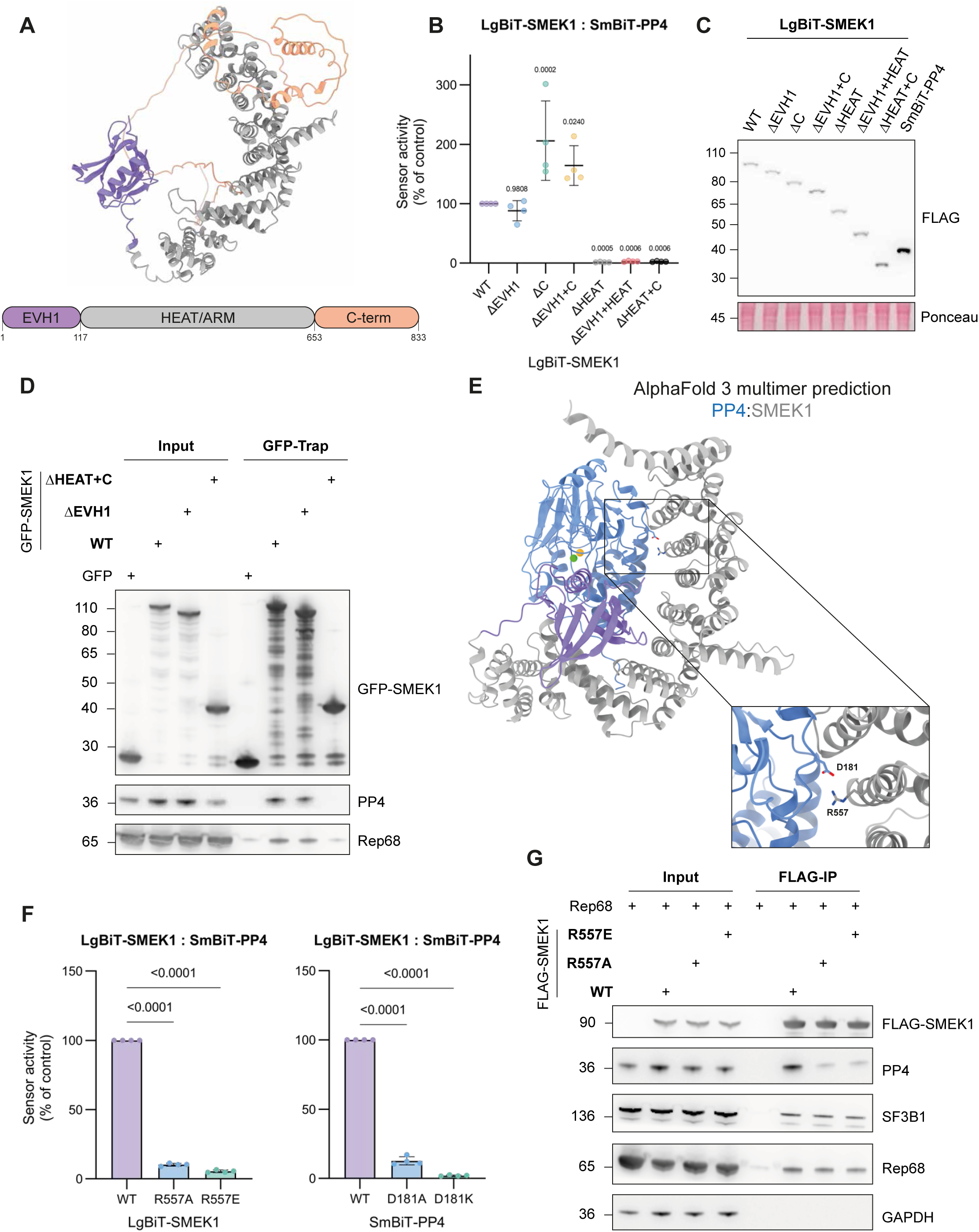
Rep68 interacts with the HEAT/Arm repeat domain of SMEK1, independently of PP4. **(A)** SMEK1 domain structure from the AlphaFold Protein Structure Database ^77,78^. The EVH1, HEAT/Arm and C-terminal domain are indicated in purple, grey and orange respectively. **(B)** LgBiT-SMEK1 deletion mutants tested for their interaction with SmBiT-PP4. Lysate-based split-luciferase assay performed by mixing HEK293T lysates from cells that separately expressed LgBiT-SMEK1 (WT or mutant) and SmBiT-PP4. Bioluminescence signal was measured after 20 minutes incubation at room temperature (end-point measurement). The data was plotted as a percentage of the LgBiT-SMEK1^WT^:SmBiT-PP4 signal (mean ± SD; n = 4 independent experiments). Statistical significance was determined by two-sided unpaired t-test. **(C)** Transient expression levels of LgBiT-SMEK1 (WT or mutants) and SmBiT-PP4 assessed via immunoblotting of the lysates used in **B**. **(D)** GFP-trap results of ectopically expressed GFP-SMEK1 (WT and mutants) showing loss of binding between GFP-SMEK1^ΔHEAT+C^ and Rep68. **(E)** AlphaFold 3 multimer prediction of the PP4:SMEK1 holoenzyme complex. The identified PP4^D181^:SMEK1^R557^ salt bridge is shown as sticks in the close-up panel. PP4 active site metals, Zn^2+^ and Fe^2+^, are represented as a green and orange sphere respectively. **(F)** Lysate-based split-luciferase end-point measurements of the LgBiT-SMEK1^R557➔A/E^:SmBiT-PP4^WT^ (left panel) and LgBiT-SMEK1^WT^:SmBiT-PP4^D181➔A/K^ (right panel) mutants interaction sensors. Bioluminescence signal was measured after 20 minutes of incubation at room temperature and plotted as a percentage of the LgBiT-SMEK1^WT^:SmBiT-PP4^WT^ signal (mean ± SD; n = 4 independent experiments). Statistical significance was determined by two-sided unpaired t-test. **(G)** FLAG-IP results of ectopically expressed FLAG-SMEK1 (WT and R557 mutants) showing loss of binding between endogenous PP4 and FLAG-SMEK1^R557➔A/E^, while the interaction with ectopically expressed Rep68 is unaffected.

Since the SMEK1:Rep68 and SMEK1:PP4 interaction both involve the HEAT/Arm domain of SMEK1, we subsequently investigated if PP4 association with SMEK1 is necessary for the SMEK1:Rep68 interaction. As there are no structural models for PP4:SMEK1, we used AlphaFold 3 to predict the structure of the PP4:SMEK1 complex^44^. This resulted in five models with high confidence scores, all converging on a single structural topology of the PP4:SMEK1 complex. Using this model, we could identify a salt bridge between SMEK1^R557^ and PP4^D181^ (Figure 2E). Mutation of these residues to alanine or its opposite charge largely abolished their interaction, as observed in both the lysate-based split-luciferase and FLAG-IP assays (Figure 2F,G and S2D). Using thermal shift assays, we confirmed that loss of binding between LgBiT-SMEK1^R557A/E^ and PP4 was not due to altered protein folding of SMEK1 (Figure S2E). The PP4-binding mutants still interacted with Rep68 (Figure 2G and S2F), indicating that the interaction between Rep68 and SMEK1 is independent of PP4 within the complex. The interaction of GFP-SMEK1 with SF3B1 was not affected upon R557 mutation, delivering additional proof that the GFP-SMEK1^R557A/E^ conformation was not altered (Figure 2G).

Collectively, these data show that the Rep proteins target the HEAT/Arm domain of SMEK1, independently of PP4.

### Removal of PP4 or its regulatory subunits enhances AAV replication and gene expression

Because the AAV Rep proteins interact directly with SMEK1, we speculated that they might interfere with PP4 phosphatase activity to prevent dephosphorylation of substrates that support viral replication. To examine whether the PP4 phosphatase affects AAV replication, we generated a doxycycline-inducible shRNA-mediated PP4 knockdown (KD) HEK293T cell line. This cell line contains a stable genomic integration of a sequence encoding an shRNA that targets the 3’-UTR of *PPP4C,* under the control of a doxycycline-inducible promoter. We found that doxycycline-induced knockdown of PP4 was associated with enhanced viral protein expression levels and increased AAV replication (Figure 3A-C). Furthermore, knockdown of PP4 resulted in a marked increase in KAP1 phosphorylation, while KAP1 expression levels remained unchanged.

**Figure 3.**
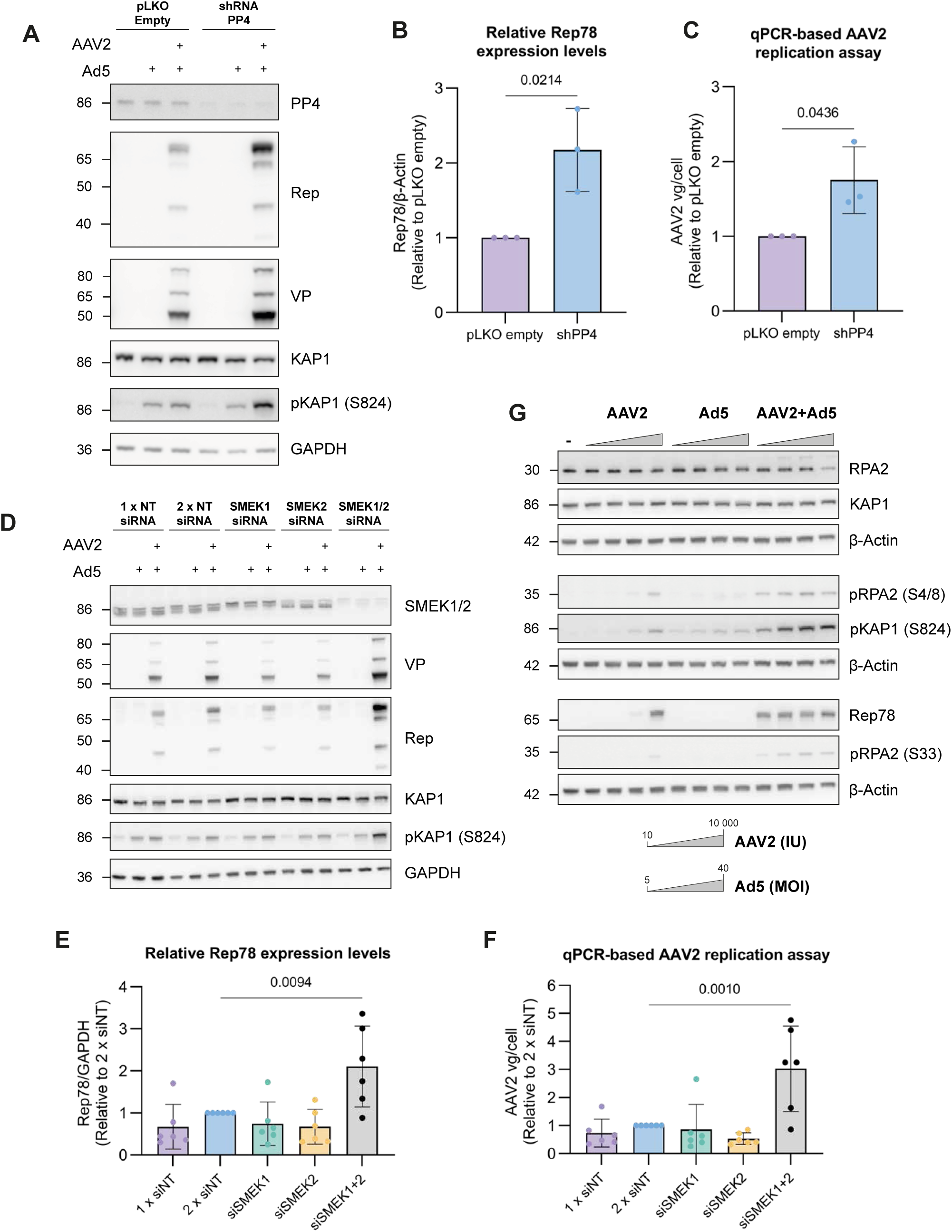
PP4:SMEK1/2 holoenzyme is a repressor of AAV replication. **(A-C)** AAV replication assay in shRNA-mediated PP4C knockdown HEK293T cell lines. Knockdown of PP4C was induced by treating the cells with 1 μg/mL doxycycline (Dox) for 72h after which they were infected with 10 IU AAV2 and a MOI 5 of Ad5. Cells were harvested 28h post-infection for analysis of PP4 knockdown and viral Rep/VP expression levels via immunoblotting in **A** and **B**, and for quantification of the viral genomes per cell (vg/cell) via qPCR in **C** (mean ± SD; n = 3 independent experiments). Statistical significance was determined by unpaired t-test. **(D-F)** AAV replication assay after siRNA-mediated depletion of SMEK1 and SMEK2. siRNA against SMEK1 and SMEK2 were transiently transfected in HEK293T. SMEK1/2 depleted cells were infected 72h post-transfected with 10 IU AAV2 and a MOI 5 of Ad5. Cells were harvested 28h post-infection and processed as in **A-C** (mean ± SD; n = 6 independent experiments). Statistical significance was determined by two-sided unpaired t-test. (**G)** Effect of lytic AAV2 infection on the phosphorylation status of KAP1^S824^, RPA2^S4/8^ and RPA2^S33^. HEK293T cells were infected with increasing IUs of AAV2 and Ad5.

Since Rep interacts directly with SMEK1, we also carried out siRNA-mediated knockdown experiments to assess the effect of SMEK1 depletion on AAV replication. Surprisingly, KD of SMEK1 alone had no discernible effect on viral replication (Figure 3D–F). Given the extended sequence and structural homology between SMEK1 and SMEK2 (Figure S3A,B), we next investigated whether simultaneous knockdown of both subunits would mimic the effect observed with PP4 knockdown. Indeed, the combined knockdown of SMEK1 and SMEK2 significantly enhanced viral replication and gene expression, indicating functional redundancy between these regulatory subunits (Figure 3D-F). This functional redundancy is consistent with their high structural similarity and is further supported by CRISPR-based dependency data from the DepMap project, which show that SMEK1 and SMEK2 are functionally co-dependent in a wide range of cell lines (https://depmap.org/portal/) (Figure S3C). In view of the structural and functional similarity between SMEK1 and SMEK2, we next examined whether Rep proteins also interact with SMEK2. GFP-trap assays using GFP-tagged SMEK1 and SMEK2 revealed robust co-precipitation of Rep68 with both SMEK proteins, indicating that Rep68 can associate with either regulatory subunit (Figure S3D).

Finally, we observed that infection of HEK293T cells with increasing AAV infection units (IU) in the presence of a fixed multiplicity of infection (MOI) of Ad5, led to a marked increase in phosphorylation of RPA2 at residues Ser4, Ser8 and Ser33 which are well-established targets of PP4 (Figure 3G) ^36^. In contrast, Ad5-only infection did not induce RPA2 phosphorylation, indicating that this hyperphosphorylation is specifically mediated by the AAV Rep proteins. These findings are in line with our previous observation of elevated KAP1^S824^ phosphorylation during lytic AAV infection (Figure 3G) ^19^.

Together, these findings demonstrate that the PP4:SMEK1 and PP4:SMEK2 holoenzymes function as negative regulators of AAV replication and gene expression. This repression is most likely alleviated through Rep-dependent inhibition of PP4 phosphatase activity, through a sustained hyperphosphorylation of PP4 substrates such as KAP1 and RPA2.

### The PP4 substrate KAP1 interacts with SMEK1 and SMEK2 via its MAPP short linear motif

To better understand how KAP1 is recruited to PP4 via its regulatory subunits, we investigated the specific interaction sites between KAP1 and SMEK1/2. PP4:SMEK1/2 substrates usually harbour a conserved FxxP/MxPP SLiM motif, which binds with low micromolar affinity to a hydrophobic groove in the EVH1 domain of SMEK1/2 ^34^. KAP1 contains a single such motif, ^423^MAPP^426^ (Figure 4A). SLiMs are typically characterized by two or three key residues located within a sequence of up to ten amino acids, often found in intrinsically disordered regions of the proteins involved in the interaction ^45^. The ^423^MAPP^426^ SLiM motif is located N-terminal to the Ser473 and Ser824 residues, which are phosphorylated by Chk2 and ATM, respectively, during the DNA damage response, leading to chromatin relaxation. Following DNA repair, Ser473 and Ser824 residues are dephosphorylated by PP4 ^33,46^. The AlphaFold-predicted structure of KAP1 suggests that its ^423^MAPP^426^ motif resides within an intrinsically disordered region (Figure 4A). Alanine substitution of this motif strongly reduced its interaction with LgBiT-SMEK1 in lysate-based split-luciferase assays (Figure 4B and S4A). Additionally, deletion of the conserved EVH1 domain from SMEK1 decreased its interaction with KAP1 (Figure S4B), indicating that the MAPP motif of KAP1 interacts with the EVH1 motif of SMEK1.

**Figure 4.**
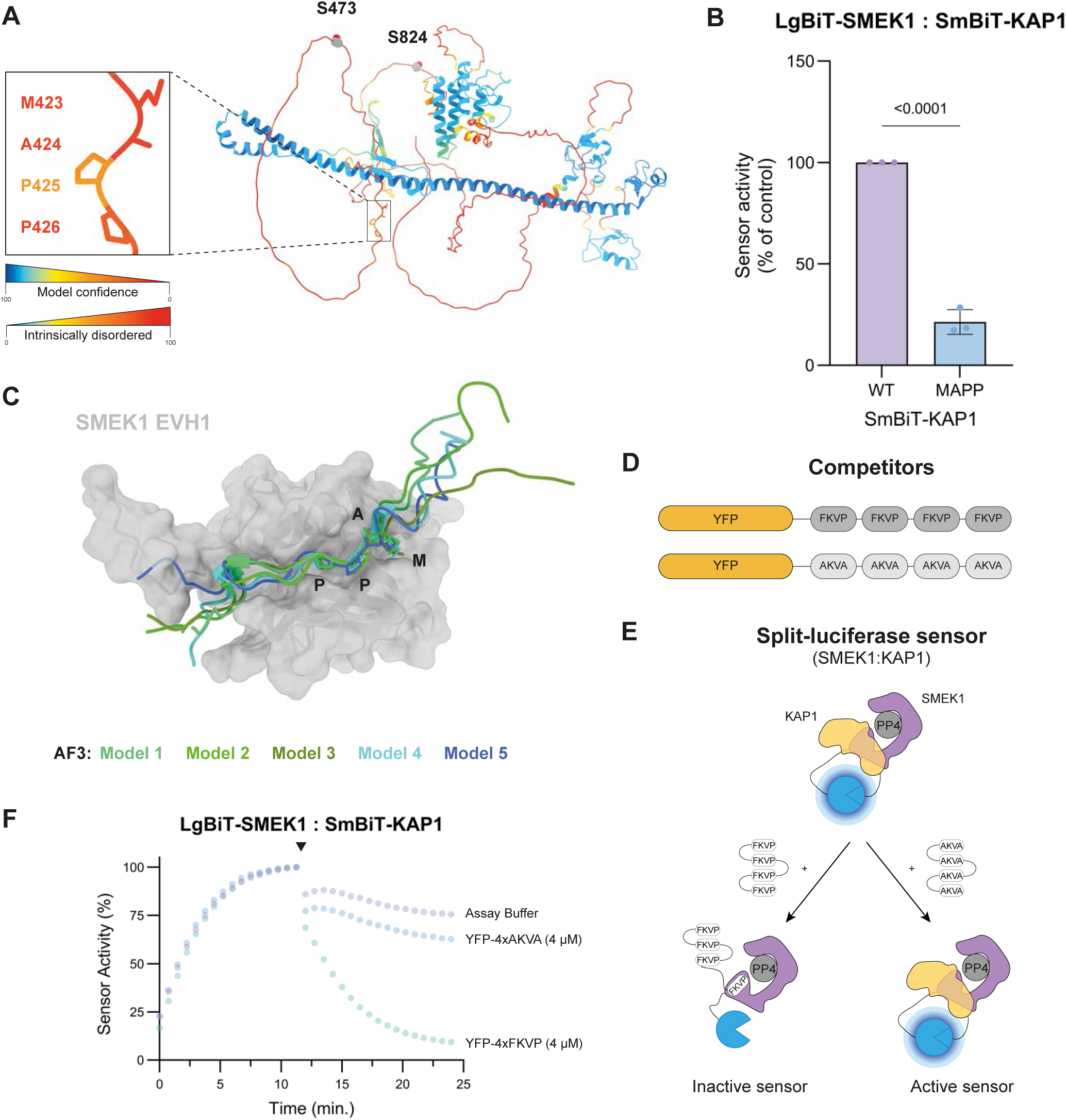
KAP1 interacts via is MAPP SLiM with the EVH1 domain of SMEK1 and SMEK2. **(A)** KAP1 (TRIM28) tertiary structure modelled via the AlphaFold Protein Structure Database. The structure is coloured according to their local model confidence. The lower the model confidence, the higher the intrinsically disordered character. MxPP SLiM (^423^MAPP^426^) highlighted as sticks in the close-up view (left panel). **(B)** Lysate-based split-luciferase assay (end-point measurement) of the LgBiT-SMEK1:SmBiT-KAP1^MAPP^ interaction sensor. Bioluminescence signal was measured after 20 minutes of incubation at room temperature (end-point measurement). The data was plotted as a percentage of the LgBiT-SMEK1^WT^:SmBiT-KAP1^WT^ signal (mean ± SD; n = 3 independent experiments). Statistical significance was determined by two-sided unpaired t-test. **(C)** AlphaFold 3 multimer prediction of the SMEK1^EVH1^ domain in complex with the KAP1^413–437^ peptide containing the MAPP SLiM. The five obtained models were structurally aligned. **(D)** Design of the YFP-4xFKVP competitor peptides. YFP-4xAKVA serves as the non-binding control peptide. **(E)** Design of the kinetic-trace experiments using the LgBiT-SMEK1:SmBiT-KAP1 interaction sensor and YFP-4xFKVP as competitor. For simplicity, the YFP-tagged competitor and control peptide are represented without YFP. **(F)** Kinetic-trace experiment showing the time-dependent association of the LgBiT-SMEK1:SmBiT-KAP1 split-luciferase sensor. The black arrow indicates the addition of the purified YFP-4xFKVP/AKVA competitor (concentration indicated in the graph). The represented data is plotted as a percentage of the S/B ratio right before the addition of the competitor. The data shown is a representative example of three independent repeats.

The interaction of the KAP1 MAPP motif with SMEK1’s EVH1 domain was further validated using AlphaFold 3 multimer predictions. Modeling of the SMEK1 EVH1 domain with a KAP1-derived peptide containing the MAPP motif yielded five high-confidence models that converged on a single topology. In all models, the peptide was positioned within the hydrophobic groove of the EVH1 domain, resembling the binding mode of endogenous PP4 substrates that engage SMEK1/2 via an FxxP SliM (Figure 4C) ^34^. Structural alignment of the SMEK1^EVH1^:KAP1^MAPP^ AlphaFold model with the published crystal structure of SMEK1^EVH1^ in complex with a model FxxP peptide revealed that both peptides aligned with a RMSD of 0.539 (Figure S4C) ^34^.

We next utilized the EVH1-binding peptide (SLPFT<u>FKVP</u>APPPSLPPS) described by Ueki et al. for competition studies ^34^. We fused four copies of this peptide to YFP (YFP-4xFKVP), as it was previously shown that this multivalency significantly improves the binding affinity of SLiMs, enhancing its potency as a competitive inhibitor of SMEK1/2 substrate interactions ^47^. The YFP-4xFKVP competitor peptide effectively disrupted the SMEK1:KAP1 interaction in lysate-based split-luciferase assays (Figure 4D-F and S4D). A similar effect was observed for the LgBiT-SMEK2:SmBiT-KAP1 interaction sensor (Figure S4E). In contrast, a control peptide (YFP-4xAKVA) barely affected the SMEK1/2:KAP1 interaction.

In summary, we were able to disrupt the SMEK1:KAP1 interaction by mutating the ^423^MAPP^426^ motif in KAP1, deleting the EVH1 domain of SMEK1 or through competitive displacement by adding the YFP-4xFKVP competitor, demonstrating that KAP1 binds the EVH1 domain of SMEK1 at the canonical ‘FxxP‘-substrate interaction site via its ^423^MAPP^426^ SLiM.

### Rep68 prevents the recruitment of PP4 substrates via SMEK1/2

The observation that lytic AAV infection, and the associated expression of Rep proteins, results in increased phosphorylation of PP4 substrates, led us to speculate that Rep proteins inhibit PP4 holoenzymes. One possible mechanism is through interference with substrate recruitment. We performed various lysate-based split-luciferase assays to test whether Rep68 interferes with substrate recruitment of the PP4:SMEK1 holoenzyme. First, we observed that the purified KAP1 competes with Rep68-SmBiT for binding to LgBiT-SMEK1 (Figure 5A). Conversely, addition of purified Rep68 also competed with SmBiT-KAP1 binding to LgBiT-SMEK1 (Figure 5B). This indicates that both KAP1 and Rep68 share overlapping interaction sites on SMEK1.

**Figure 5.**
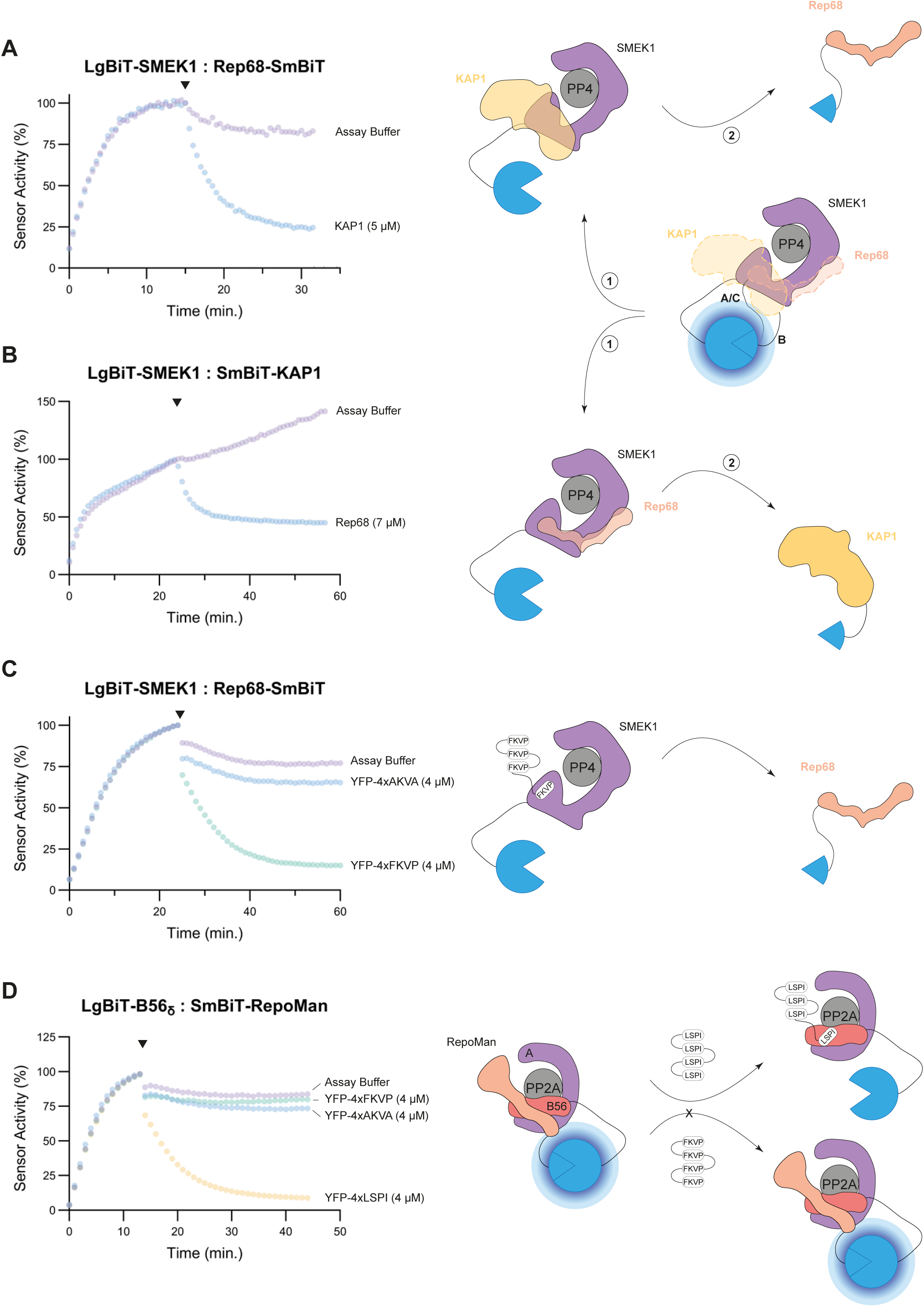
AAV Rep68 interferes with the recruitment of PP4:SMEK1 substrates. **(A)** Kinetic-trace experiment showing the time-dependent association of the LgBiT-SMEK1:Rep68-SmBiT split-luciferase sensor. The black arrow indicates the addition of the purified, untagged KAP1 competitor (concentration indicated in the graph). The represented data is plotted as a percentage of the S/B ratio right before the addition of the competitor. The data shown is a representative example of three independent repeats. **(B)** Same experiment as in **A**, but with the LgBiT-SMEK1:SmBiT-KAP1 split-luciferase sensor and purified, untagged Rep68 as competitor. **(C)** Analogous experiment as in **A**, but with YFP-4xFKVP/AKVA as competitor. **(D)** Kinetic-trace experiment showing the time-dependent association of the LgBiT-B56_8_:SmBiT-RepoMan split-luciferase sensor. The black arrow indicates the addition of the purified YFP-competitor peptides (concentration indicated in the graph). The represented data is plotted as a percentage of the S/B ratio right before the addition of the competitor. The data shown is a representative example of three independent repeats.

To further validate the competition between KAP1 and Rep68 for binding to SMEK1, we used the YFP-4xFKVP peptide as a competitor in our lysate-based split-luciferase assays (Figure 5C). Addition of the YFP-4xFKVP competitor peptide to the LgBiT-SMEK1:Rep68-SmBiT interaction sensors resulted in dissociation of the protein complex, while the control peptide had no significant effect on the interaction (Figure 5C). This confirms the data obtained in figure S2C showing that Rep68 interacts with the EVH1 domain of SMEK1, and thereby probably interferes with substrate recruitment. However, previous pull-down and split-luciferase experiments (Figure 2D and S2C) showed that the HEAT/Arm repeat domain of SMEK1 is the main determinant for the SMEK1:Rep68 interaction. The YFP-4xFKVP peptide produced similar effects when added to the LgBiT-SMEK2:Rep68-SmBiT interaction sensor (Figure S5A).

To confirm the specificity of the effect of the FKVP peptide on EVH1-mediated interactions, we performed a control experiment using the LgBiT-tagged B56_δ_ regulatory subunit of PP2A and SmBiT-RepoMan, a known PP2A substrate. RepoMan recruitment to PP2A via B56δ depends on the LSPI SLiM and thus should not be affected by the YFP-4xFKVP peptide. The PP4-specific SLiM peptide showed no effect on the B56_δ_:RepoMan interaction, whereas the YFP-4xLSPI peptide almost completely dissociated the interaction sensor (Figure 5D), confirming the selectivity of the YFP-4xFKVP competitor peptide for EVH1 binding substrates ^34^.

In conclusion, these data show that Rep68 and KAP1 may share overlapping binding sites on SMEK1, or that steric hinderance upon SMEK1 binding of the viral protein prevents the binding of the other PP4 substrates. Furthermore, we demonstrate that the interaction of both proteins with SMEK1 are at least in part mediated by the FxxP docking site on the EVH1 domain of SMEK1/2.

### The NIPP1 regulatory subunit of PP1 interacts with the SMEK1 regulatory subunit of PP4

We previously reported that AAV Rep proteins also target the PP1:NIPP1 holoenzyme complex to promote viral replication and transcription ^19^. This prompted us to explore whether PP1:NIPP1 and PP4:SMEK1 are part of the same macromolecular complex. For this reason, we performed FLAG-IP experiments from cells that were transiently transfected with FLAG-tagged full-length NIPP1^WT^, NIPP1^Δ1–22^ lacking the first 22 amino acids known as the FHA-inhibitory domain (FID) ^40^ or NIPP1^Δ1-142^ lacking the FID and substrate-binding FHA domain (Figure 6A). As expected, binding of Rep68 and the substrates KAP1 and SF3B1 to NIPP1, depend on the N-terminal FHA domain of NIPP1 ^19,37^. Surprisingly, we also observed an association between endogenous SMEK1 and FLAG-NIPP1^WT^. This association was not affected by deletion of the FID domain (FLAG-NIPP1^Δ1–22^) but was abolished by the additional deletion of the FHA domain (NIPP1^Δ1-142^) (Figure 6A). This was confirmed by a GFP-trap of GFP-SMEK1 from cells ectopically expressing either FLAG-tagged wild-type or NIPP1^Δ1-142^ (Figure 6B). Additional GFP-trap experiments with truncated mutants of GFP-SMEK1 indicated that the HEAT/Arm domain of SMEK1 is important for the association with endogenous NIPP1 (Figure S6A).

**Figure 6.**
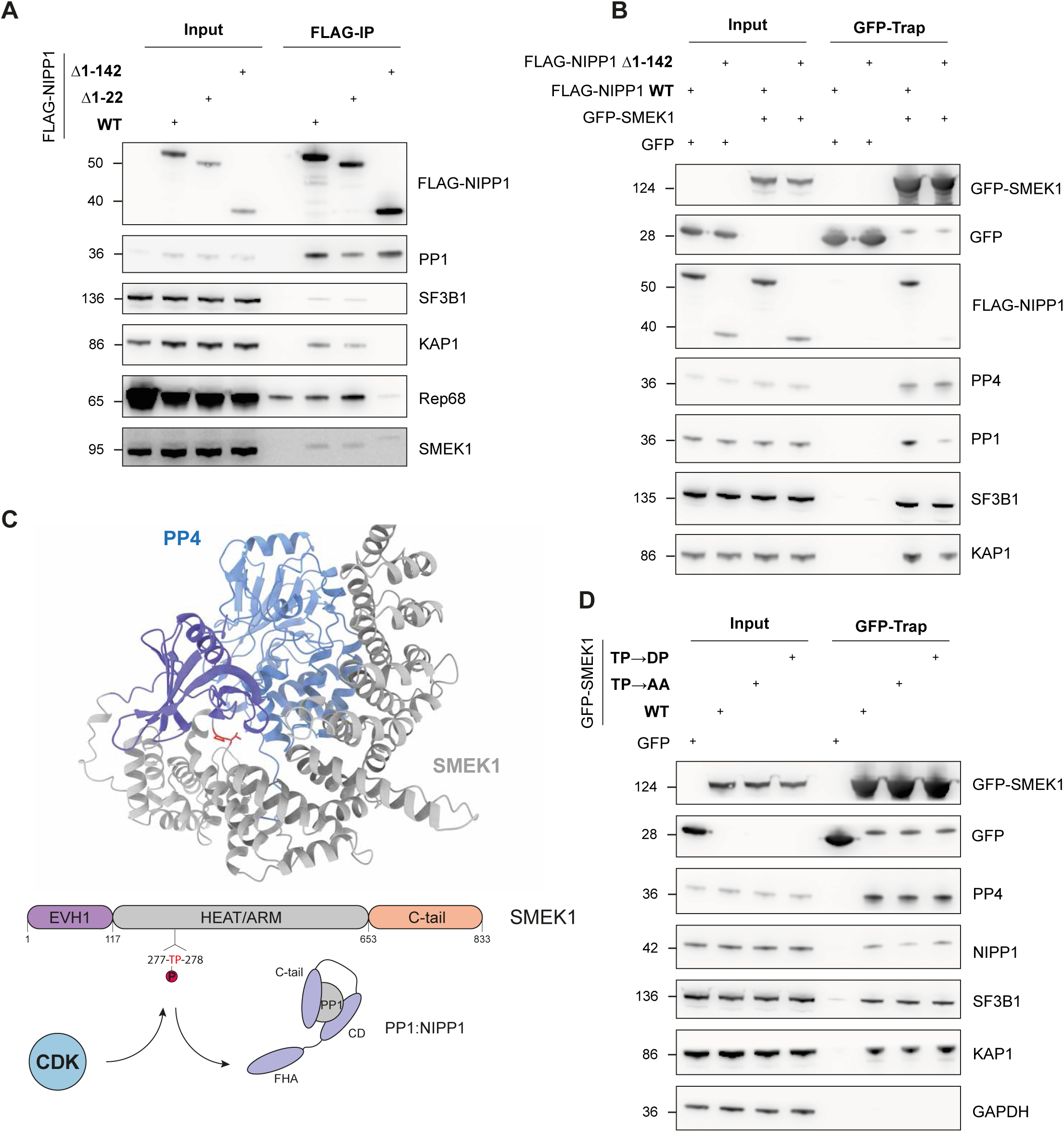
Rep interacts with the putative PP1:NIPP1-PP4:SMEK1 multifunctional phosphatase complex. **(A)** FLAG-IP of FLAG-NIPP1 (WT, Δ1-22 and Δ1-142) transiently expressed in HEK293T. **(B)** GFP-trap of GFP-SMEK1 to investigate the co-precipitation of FLAG-NIPP1 (WT and Δ1-142) transiently expressed in HEK293T cells. **(C)** AlphaFIold 3 model of the PP4:SMEK1 complex, with the SMEK1^TP^ dipeptide motif highlighted in red as sticks. CDK kinases might phosphorylate SMEK1^T277^, thereby increasing the affinity of SMEK1 for NIPP1^FHA^ binding. **(D)** GFP-trap of GFP-SMEK1 (TP ➔ AA or DP) to investigate the effect on co-precipitation of endogenous NIPP1.

NIPP1 recruits substrates for associated PP1 through its FHA domain, which recognizes a phosphorylated threonine-proline (pTP) motif on the substrate, where the threonine is phosphorylated by cyclin-dependent kinases (CDKs). Known PP1:NIPP1 substrates that interact via this mechanism include SF3B1, CDC5L, MELK, EZH2 ^37,48,49^. We identified a TP-dipeptide motif (^277^TP^278^) in SMEK1, located in a disordered connecting loop in the HEAT/Arm domain (Figure 6C). A GFP-trap was performed using GFP-SMEK1 in which the TP-dipeptide motif was mutated to AA (phosphomutant) or DP (phosphomimetic). The TP➔AA mutant of SMEK1 resulted in a slightly decreased association with NIPP1, while no strengthened binding of the TP➔DP mutant was observed (Figure 6D). This might be due to the unknown extent of phosphorylation in the WT construct and uncertainty whether mutation to aspartic acid effectively mimicks phosphorylation on this site in the TP➔DP mutant. Possibly, the bulkiness of a phosphate group is required to support proper interaction with the FHA domain of NIPP1.

Together with our previous observations ^19^, the above data indicate that PP1:NIPP1 and PP4:SMEK1 are components of the same macromolecular complex, which are targeted by AAV Rep proteins. The interaction between SMEK1 and the NIPP1 FHA domain is likely phosphorylation-independent.

## Discussion

Here, we provide for the first time evidence that a viral protein directly engages the PP4:SMEK1/2 phosphatase holoenzyme. Our findings reveal that this complex acts as a negative regulator of AAV replication and gene expression, and that its repressive activity is antagonized by the Rep proteins through a novel viral substrate-recruitment interference mechanism that serves to retain PP4 substrates in a hyperphosphorylated state (Figure 7).

**Figure 7.**
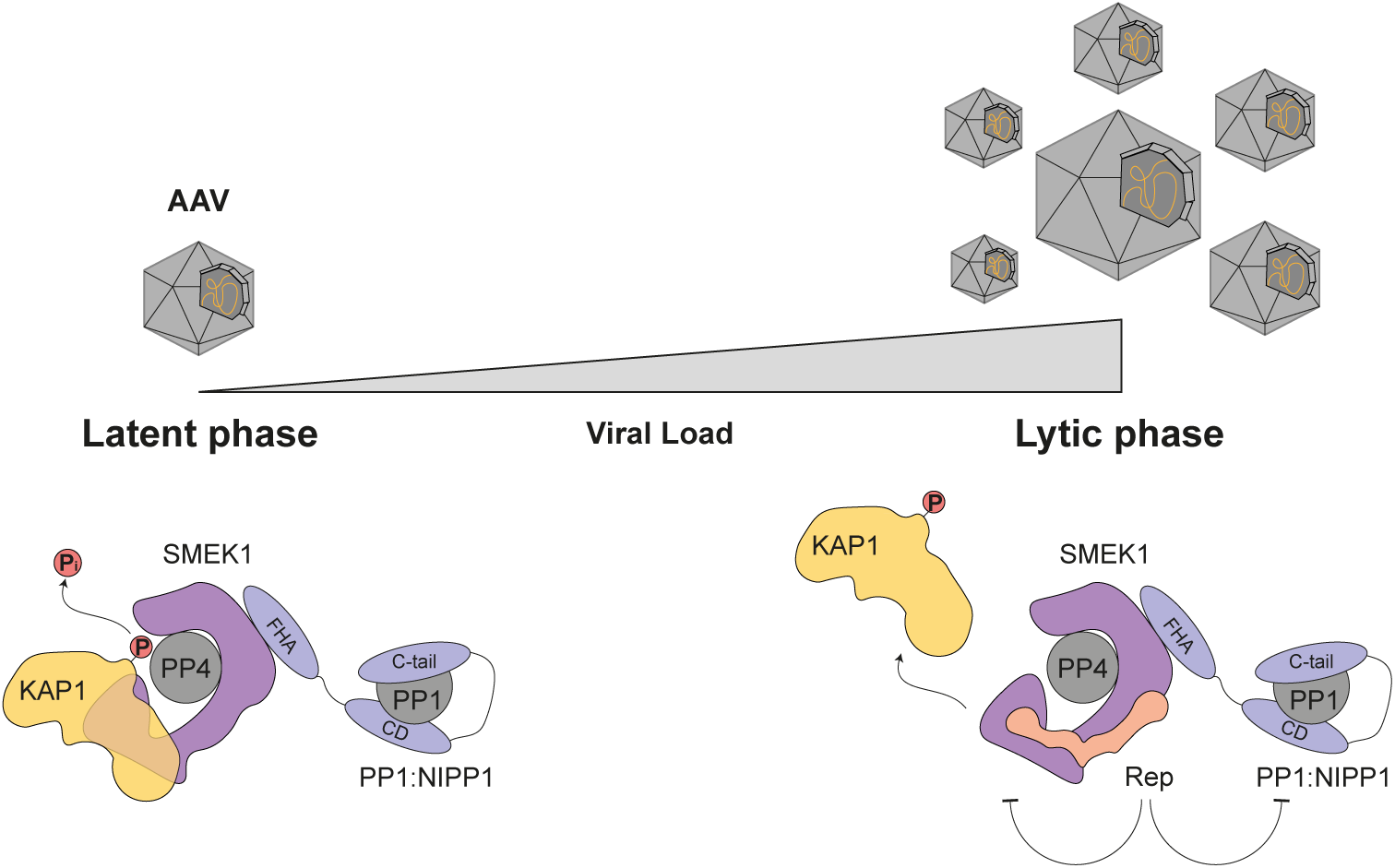
Model of the Rep-mediated substrate interference mechanism. During a latent AAV infection, in the absence of Rep expression, PP4:SMEK<u>1</u> holoenzymes are active. They actively dephosphorylate substrates that are involved in the AAV life cycle such as KAP1 and keep the AAV genome silenced. The switch from the latent to the lytic phase of the viral life cycle, upon superinfection with a helper virus, causes the expression of the viral Rep proteins, hereby boosting viral replication. The Rep proteins, together with viral replication, elicit a DNA damage response (DDR), hyperphosphorylating DDR-related proteins such as KAP1 and RPA2. The Rep proteins interact with the PP4:SMEK1 and PP1:NIPP1 holoenzymes, resulting in inhibition of PP4/PP1 phosphatase activity through interference with the recruitment of PP4:SMEK1 holoenzyme substrates. This inhibition maintains substrate hyperphosphorylation, thereby preventing repression of AAV gene expression and replication.

Hijacking of host phosphatases by viruses is a well-known strategy to modulate cellular signalling in favour of viral infectivity. Among these, the Ser/Thr phosphatases PP1 and PP2A have been extensively characterized as viral targets ^18,50^. Depending on the context, phosphatase activity may either support or hinder viral replication. Our group has previously shown that AAV Rep proteins modulate PP1 activity by interacting with the PP1:NIPP1 holoenzyme complex, ultimately leading to hyperphosphorylation of KAP1^S824^, associated with chromatin decondensation during the lytic phase of the AAV life cycle ^19^. Interestingly, PP4 has also been implicated in KAP1^S824^ dephosphorylation, suggesting that this phosphatase may serve a redundant or complementary role in this context.

SMEK1 (PP4R3A) was identified as a candidate interactor of AAV Rep proteins ^19^, prompting us to examine whether Rep proteins also target PP4 holoenzymes. To our knowledge, no prior study has reported a viral protein interacting with the trimeric PP4:PP4R2:SMEK1/2 complex. While the MCPyV small-T antigen is known to associate with PP4:PP4R1 and to modulate NF-κB signalling ^51,52^, our study characterizes a novel interaction between Rep proteins and PP4:SMEK1/2. We have found that the AAA^+^-ATPase/helicase domain, conserved across all four Rep isoforms, mainly interacts with the HEAT/Arm domain of SMEK1, a region previously shown to coordinate the PP4 catalytic subunit (Figure S1A and S2C) ^42^. AlphaFold3 modelling of the PP4:SMEK1 complex further supports this interaction, identifying a single salt bridge (PP4^D181^:SMEK1^R557^) as essential for complex formation (Figure 2E-G).

Substrates of the PP4:SMEK1/2 holoenzyme typically harbour a FxxP or MxPP SLiM within a disordered region that binds with low micromolar affinity in a conserved hydrophobic groove of the SMEK1 EVH1 domain ^34^. Based on this knowledge, we identified a similar motif (^423^MAPP^426^) in KAP1 (Figure 4A). Mutation of this motif or deletion of the EVH1 domain significantly weakened the interaction of KAP1 with SMEK1, although residual binding suggests additional contact sites. AlphaFold3 docking of the KAP1 MAPP motif on EVH1 yields five high-confidence models that all converge on a similar binding topology as observed in the EVH1:FxxP crystal structure (Figure 4C and S4C) ^34^.

Intriguingly, the AAV Rep proteins compete with endogenous, FxxP/MxPP-containing PP4:SMEK1/2 substrates such as KAP1 for SMEK1 binding, thereby blocking substrate recruitment. This substrate-recruitment inhibition mechanism was further validated in competition experiments with a synthetic validated FxxP-peptide (FKVP). These assays clearly demonstrate that the Rep proteins interfere with the recruitment of FxxP/MxPP-containing substrates. Further research is needed to identify the residues in Rep that are involved in the interaction with SMEK1 and to examine whether Rep proteins also function as a substrate for PP4-mediated dephosphorylation ^53,54^. In this respect, it is worth noting that Rep proteins are post-translationally modified by phosphorylation ^53^. Dephosphorylation of Rep proteins was suggested to be mediated by PP2A ^54^, but the underlying experiments are not conclusive as they were largely based on the use of okadaic acid (100 nM), which is not specific for PP2A but also inhibits PP4 ^55^.

Similar to the AAV Rep proteins that disrupt the interaction between PP4:SMEK1/2 and their FxxP/MxPP-containing substrates, certain viral proteins have been shown to target the PP2A:B56 holoenzyme through a conserved LxxIxE motif ^56,57^. The HTLV-1 integrase (IN), as part of the larger intasome complex, interacts with B56γ via a LxxIxE motif ^58,59^. However, the precise function of this interaction remains unknown. One hypothesis is that the intasome recruits the PP2A: B56γ holoenzyme to HTLV-1 integration sites where it may dephosphorylate transcription factors and/or chromatin modifiers to facilitate viral integration or modulate the local chromatin environment. The Ebola virus nucleoprotein (NP) recruits the viral VP30 protein to PP2A for dephosphorylation by interacting with B56 through a conserved LxxIxE motif. Dephosphorylation of VP30 switches the balance from viral replication to transcription ^15^. While both Ebola virus and HTLV-1 benefit from PP2A activity, AAV takes a different approach and inhibits PP4 function to support AAV replication and gene expression. The Rep-mediated substrate-recruitment interference described here represents a novel mechanism within the broader theme of viral targeting of PPP-type phosphatases. One hypothesis that is worth exploring is that Rep proteins inhibit substrate recruitment by functioning as an unfair competitor. During unfair competition, a phosphorylated substrate will bind its target phosphatase holoenzyme in order to be dephosphorylated. However, the dephosphorylation rates are extremely slow, hereby blocking the recruitment and dephosphorylation of other substrates. Such inhibition mechanism also applies to the inhibition of PP1:MYPT1 and PP2A:B55 by CPI-17 and endosulfine, respectively ^60,61^.

Further investigation is needed to unravel the consequence of Rep expression for the global human phosphoproteome and for specific PP4:PP4R2:SMEK1/2 substrates (e.g. γH2AX, DBC1 and 53BP1) ^35,62,63^. It is possible that binding of Rep with the HEAT/Arm and EVH1 domain of SMEK1 does not completely block the dephosphorylation of all PP4:PP4R2:SMEK1 substrates if they are recruited by an FxxP/MxPP-independent mechanism. Likewise, some PP2A:B56 substrates are recruited by an LxxIxE-independent mechanism ^64^. In the case of HTLV-1 integrase binding to B56γ via the LxxIxE motif, dephosphorylation of certain PP2A:B56γ substrates likely still proceeds, even when the LxxIxE-binding groove is occupied ^58,59^. A similar mechanism may apply to PP4 inhibition by AAV Rep proteins. A complete inhibition of PP4 by Rep proteins, through occlusion of the PP4 active site, is unlikely as this is known to be lethal, as evidenced by DepMap gene depletion screens (https://depmap.org/portal/). This highlights a critical evolutionary constraint: viruses such as AAV must rewire, rather than abolish, host phosphatase activity to avoid prematurely triggering cell death, which would be detrimental to viral replication.

Beside KAP1^S824^ hyperphosphorylation we also observe RPA2^S4/8/33^ hyper-phosphorylation, as previously described for AAV ^65^. It has been shown that the heterotrimeric RPA complex co-localizes with the Rep proteins in viral replication centres (VRCs) ^66^. Furthermore, RPA belongs to a minimal set of proteins that are needed to support AAV replication *in vitro* ^67,68^. Similar to what is observed for the autonomous parvovirus Minute virus of Mice (MVM), incoming latent AAV viral genomes can sequester substantial fractions of the nuclear RPA pool, inducing replication stress and ATR-mediated signalling ^69,70^. In addition, a fraction of RPA2 bound to the latent AAV genome appears to be phosphorylated at Ser8 ^70^. Our data suggest that hyperphosphorylated RPA2 is also associated with actively replicating AAV genomes. Whether this phosphorylation event, similar to what has been observed for KAP1, contributes to chromatin remodeling of the AAV genome remains an open question.

Remarkably, our data also revealed that SMEK1 interacts with NIPP1 and one of its substrates SF3B1 (Figure 1B and 6B), indicating that SMEK1 may function as a scaffold for the assembly of a novel multifunctional PP1:PP4 complex. This multifunctional PP1:PP4 complex may be part of the larger spliceosomal assembly, as high-throughput interaction studies reveal extensive associations between SMEK1 and multiple spliceosomal components, including SF3B1 ^71^. Several multifunctional protein complexes with phosphatases have been described before. Protein kinase A-anchoring proteins (AKAPs) bind with PKA and other kinases, thereby targeting these kinases to their substrates ^72^. Phosphatases like PP2A and PP2B can be part of such complexes to dynamically antagonize kinase signalling ^73–75^. Furthermore, one study has reported on a PP1:PP2A phosphatase complex in yeast, being essential for the re-activation of both PP1 and PP2A in late mitosis ^76^. Here, we provide evidence for the existence of a PP1:PP4 complex that involves the HEAT/Arm domain of SMEK1 and the FHA domain of NIPP1 (Figure 7). Substrates of PP1:NIPP1 often interact via a phosphorylated TP dipeptide motif ^37,48,49^. However, we found no evidence that SMEK1/2 binds to NIPP1 as substrate, indicating the SMEK1 coordinates the function of both associated PP4 and PP1. Given that both PP1:NIPP1 and PP4:SMEK1 are implicated in KAP1^S824^ dephosphorylation, this raises the intriguing possibility that Rep proteins hamper the dephosphorylation of KAP1, and possibly other substrates, by both PP1 and PP4. (Figure 7).

In conclusion, our findings reveal a novel mechanism of PPP-type phosphatase inhibition via viral-mediated substrate-recruitment interference. Furthermore, we demonstrate that the transcriptional co-repressor KAP1 primarily interacts with the EVH1 domain of SMEK1 via its MAPP SLiM. Notably, hyperphosphorylation of several repressive proteins, including KAP1 and RPA2, emerges as a recurring feature during lytic AAV infection and Rep expression. Through Rep-mediated inhibition of PP4 described here, we further illustrate the sophisticated strategy employed by Rep proteins to sustain the hyperphosphorylation of key PP4 substrates. This prevents the activation of host repressive mechanisms that would otherwise hinder viral replication, while simultaneously preserving cellular pathways that are advantageous to the viral life cycle. These insights extend our understanding of how AAV hijacks host regulatory networks and may inform strategies to enhance the efficacy and manufacturability of recombinant AAV (rAAV) vectors. Further studies will be required to determine whether phosphatases and their substrates also play a role in regulating therapeutic gene expression or contribute to the complex host-cell interactions underlying rAAV production. Such knowledge could uncover new molecular targets for engineering more potent vectors and optimizing clinical gene therapy manufacturing platforms. In parallel, the growing interest in targeting phosphatases for therapeutic purposes highlights the broader relevance of this work, which may offer new insights into how phosphatase activity can be modulated to treat diseases such as cancer.

## Materials and Methods

### Protein expression and purification

His-Rep68, His-KAP1, His-LgBiT-SMEK1 (WT and deletion mutants), His-Rep68-SmBiT, His-SmBiT-KAP1, His-YFP-4xFKVP, His-YFP-4xAKVA and His-YFP-4xLSPI were expressed from the pET16b vector in BL21 Gold *E. coli* cells. Cells were grown in Luria Bertani broth, supplemented with 100 μg/mL ampicillin, at 37 °C until the optical density at 600 nm reached 0.6-0.7. Next, recombinant protein expression was induced by adding 1 mM IPTG to the cell suspension. Protein expression was induced overnight at 18 °C. Cells were pelleted by centrifugation (6000 x g, 10 min., 4 °C) and resuspended in lysis buffer containing 50 mM Tris at pH 7.9, 0.5 NaCl, 10 % glycerol, 0.5 % Triton X-100, 1 mM phenylmethylsulfonyl fluoride (PMSF), 1 mM benzamidine and 0.5 μg/mL leupeptin (LPP). Cell suspensions were subjected to one freeze-thaw cycle, followed by incubation with 10 U benzonase and 0.5 mg lysozymes per mL cell lysate for 20 minutes at room temperature. After the benzonase and lysozyme treatment, the lysates were sonicated for 15 minutes with a Diagenode Bioruptor sonicator (15 min., 15 sec. ON, 15 sec. OFF). The debris was spun down (15 000 x g, 20 min., 4 °C) and cleared lysates were incubated with Ni^2+^-Sepharose beads for 1 h at 4 °C. The beads were washed two times with lysis buffer and afterwards with buffer containing 50 mM Tris at pH 7.9, 0.5 M NaCl and imidazole concentrations of either 20 or 60 mM. Proteins were eluted from the beads by applying the same Tris-based buffer to the beads containing 0.4 M imidazole. Eluted proteins were dialyzed to a buffer containing 50 mM Tris at pH 7.5 and 150 mM NaCl. Purity of the purified proteins was confirmed by SDS-PAGE and Coomassie staining. Concentration of the purified proteins was determined via NanoDrop spectrophotometer measurements of the absorbance at 280 nm.

### Cell culture and treatments

HEK293T cells were cultured in Dulbecco’s modified Eagle’s medium (Gibco™) with 4.5 mg glucose/mL, supplemented with 10 % fetal calf serum (Sigma-Aldrich), 100 units penicillin (P)/mL and 100 μg streptomycin (S)/mL (Gibco™). Cells were transiently transfected with plasmid DNA using polyethylenimine (PEI) MAX, with a 1:4 (DNA:PEI MAX) ratio diluted in DMEM without serum and antibiotics. Cells were harvested 48 h post-transfection. Transient gene knockdowns of SMEK1 and SMEK2 were obtained through transient transfection of freshly seeded HEK293T cells, in DMEM with FCS but without P/S, with JetPrime® (Polyplus) and the desired ON-TARGET plus siRNA SMARTpool mix from Horizon discovery (see supplementary table). Cells were transfected with 35 nM siRNA according to the JetPrime® transfection protocol for siRNA. Cells transiently transfected with ON-TARGETplus non-targeting control pool served as a control. Optimal gene knockdown was achieved 72 h post-infection.

### Generation of shRNA PP4C knockdown cell line and treatment

All primers and the shRNA oligos used in this section are detailed in supplementary table. Oligos of the top and bottom strand of the shRNA targeting the 3’-UTR of the PPP4C gene were ordered from Integrated DNA Technologies (IDT) and annealed to each other using the thermocycler (Biometra TOne) starting at 95 °C for 5 minutes, followed by a steady decrease in temperature during 45 minutes until 20 °C was reached. The annealed oligos were cloned in the AgeI and EcoRI linearized Tet-pLKO-puro backbone using NEBuilder® HiFi DNA assembly by mixing equimolar ratios of both fragments. Correct ligation of the shRNA oligo in the Tet-On-pLKO backbone was verified through Sanger sequencing via LGC Genomics (Berlin, Germany). The Tet-On-pLKO-PP4C-3’-UTR plasmid was then used to produce lentiviral particles carrying the PP4C shRNA targeting the 3’-UTR. 1*10^7^ HEK293T cells were seeded in a 10 cm dish followed by transient transfection with 4 µg pRSv REV, 2.5 µg pMD2 VsVg and 4 µg pMDLgag/pRRE #54 helper plasmids and 5 µg of the Tet-On-pLKO-PP4C-3’-UTR plasmid (detailed in supplementary table) using PEI Max. The cell culture medium was filtered three days post-transfection through a 0.22 μm filter and aliquots of the virus stored at −80 °C until further use. Lentiviral titters were determined by ELISA. Transduction of HEK293T cells at a multiplicity of infection (MOI) of 25 was performed by seeding the cells at a confluency of 20 % in a 10 cm dish already containing the lentiviral particles pre-mixed with polybrene. Medium was changed 24 h post-transduction. Transduced cells were split and subjected to puromycin selection (1 μg/mL) three days post-transduction. The HEK293T shRNA-PP4C-3’-UTR cell line was stored in liquid nitrogen after four passages in the presence of puromycin. PP4C knockdown was induced by treating the cells with 1 μg/mL doxycycline for at least 72 h. Puromycin selection (1 μg/mL) was maintained during the experiments.

### Immunoblotting and immunoprecipitation

Total cell lysates were prepared by harvesting the cells and resuspending them in 10 pellet volumes of total cell lysis (TCL) buffer containing 50 mM Tris at pH 7.4, 150 mM NaCl, 0.1 % sodium dodecyl sulfate (SDS), 0.5 % sodium deoxycholate, 1 % nonident P-40 (NP-40), 1 mM dithiothreitol (DTT), 1 mM PMSF, 1 mM benzamidine, 0.5 μg/mL LPP, 50 mM NaF, 10 mM β-glycerophosphate, 1 mM Na_3_VO_4_. Crude cell lysates were sonicated with a Hielscher handheld probe sonicator (5 sec., cycle 1, amplitude 70 %) and spun down to pellet debris (15 000 x g, 10 min., 4 °C). Cleared lysates were boiled in SDS sample buffer and analyzed by SDS-PAGE using Bis-Tris NuPAGE^®^ 4-12 % gels. Proteins were blotted on a nitrocellulose membrane for 2 hours at 40 V in buffer containing 50 mM Tris and 50 mM boric acid, followed by Ponceau S staining to assess equal loading in all lanes. Next, the membranes were incubated overnight with primary antibody diluted in 5 % milk/TBS-Tween20. Next, the membranes were washed three times with TBS-T, followed by 1 h incubation with HRP-coupled secondary antibody at room temperature.

Immunoblots were eventually analysed using an ImageQuant LAS 4000 and ECL reagent (PerkinElmer). Lysates for immunoprecipitations were prepared from cells from two 15 cm dishes, lysed in Co-IP buffer composed of 10 mM HEPES at pH 7, 150 mM NaCl, 6 mM MgCl_2_, 0.5 % NP-40, 10 % glycerol, 2 mM DTT, 1 mM PMSF, 1 mM benzamidine, 0.5 μg/mL LPP. Crude lysates were sonicated with a Diagenode bioruptor sonicator (10 min., 15 sec. ON, 15 sec. OFF), followed by centrifugation (1000 x g, 10 min., 4 °C). 60 μL of the cleared lysates was kept as input samples and boiled with sample buffer. The remaining lysate was either incubated with 30 μL of home-made anti-EGFP nanobodies covalently coupled with agarose beads for GFP-traps, or 30 μL anti-FLAG^®^ M2 affinity gel resin for FLAG-IPs. Lysates were incubated with the beads for 2h at 4 °C, before being washed with lysis buffer. The anti-EGFP nanobody beads were directly boiled in sample buffer, while the anti-FLAG^®^ M2 affinity gel resin was incubated with 3X-FLAG peptide for 1 h at 4 °C to elute the proteins. Eluted proteins were then boiled in sample buffer.

### Split-luciferase assays

For the split-luciferase assays, protein-protein interaction sensors were made based on the NanoBiT technology (Promega) and the rationale described in Claes and Bollen, 2023 ^38,41^. In most of our assays, SMEK1 was tagged with the LgBiT-tag, by cloning SMEK1 cDNA into a vector containing the LgBiT-tag. Other proteins were fused with the SmBiT-tag. Flexible linkers, ranging from 10-20 amino acids, were included between the tags and the protein.

7 μg of the plasmids expressing the LgBiT- and SmBiT-tagged proteins were transiently transfected in a 15 cm dish containing HEK293T cells at a confluency of 25 % using PEI Max as the transfection agent. Medium was changed for fresh medium 24 h post-transfection. 48 h post-transfection, cells were harvested and cell pellets were resuspended in 10 pellet volumes of assay buffer containing 50 mM Tris at pH 7.4, 150 mM NaCl, 10 % glycerol, 0.01 % saponin, 0.5 mM EDTA, 1 mM PMSF, 1 mM benzamidine, 0.5 μg/mL LPP and 1 mM DTT. Cell suspensions were freeze-thawed once, after which the lysates were cleared (20 000 x g, 5 min., 4 °C). These cell lysates were further used for the lysate-based split-luciferase assays, as previous described^38,41^.

In general, lysate-based split-luciferase assays were carried out in white, non-binding 384-well plates (Greiner) (deep-well or high-base). Cell lysates containing the SmBiT-tagged proteins were pipetted in the wells before cell lysate containing the LgBiT-tagged protein supplemented with the furimazine substrate (50 μM final) was added. For end-point measurements, the plate was analysed after 15 to 20 minutes incubation at room temperature, while for kinetic measurements, the plate was read out every 10 to 20 seconds, until equilibrium was reached. All measurements were carried out at room temperature. For each measurement, a condition with the LgBiT-tagged cell lysate alone (addition of assay buffer instead of SmBiT-tagged cell lysate) was added. This accounted for the background signal, necessary to calculate the signal-to-background (S/B) ratio. Bioluminescence signal was measured using a Spark^®^ multimode microplate reader (Tecan Life Sciences). For competition assays, both LgBiT- and SmBiT-tagged cell lysates were mixed in a 384-well plate, after which the signal was continuously monitored every 10-20 seconds, until an equilibrium was reached. At this point, untagged competitor protein or assay buffer as a control was added to the wells, and the effect on the signal was monitored in function of time.

For split-luciferase assays with purified interaction sensors, 500 pM LgBiT-SMEK1 was mixed with 10 nM SmBiT-KAP1 or Rep68-SmBiT, together with 50 µM furimazine. Experimental procedures for end-point and kinetic measurements are the same as described above.

### Thermal shift assays

Assays were performed to assess the stability of mutated LgBiT-SMEK1 in HEK293T cell lysate as described before. Briefly, 50 μL aliquots of cell lysates of HEK293T cells expressing LgBiT-SMEK1 WT or R557→A/E incubated for 5 minutes in a thermocycler (Biometra TOne) with a temperature gradient ranging from 45 to 69 °C. Next, 15 μL of each incubated aliquot was transferred to a white, non-binding 384-well plate (Greiner) already containing 5 μL assay buffer with 100 μM furimazine. Thermal stability of the LgBiT-tagged proteins was determined by reading out the residual LgBiT luciferase activity using the Spark^®^ multimode microplate reader (Tecan Life Sciences).

### Viral infections

HEK293T cells at a confluency of approximately 50 % were infected with 10 infection units (IU) per cell of AAV2 and/or Ad5, at a multiplicity of infection (MOI) 5. The volume of culture medium was reduced to 2/5 of the original well volume before addition of AAV2. Cells with AA2 were incubated for 2 h at 37 °C before Ad5 was added. 1 h after Ad5 addition, the original cell culture medium was restored by addition of complete DMEM medium. The cells were harvested 28 h post-infection.

### AAV-based replication assays

HEK293T cells harvested 28h post-infection with AAV2 and/or Ad5 were divided over two clean reaction tubes. Cells were pelleted and one tube was stored at −80 °C for genomic (g) DNA extraction. The other cell pellet was lysed in total cell lysis (TCL) buffer containing 50 mM Tris at pH 7.4, 150 mM NaCl, 0.1 % sodium dodecyl sulfate (SDS), 0.5 % sodium deoxycholate, 1 % nonident P-40 (NP-40), 1 mM dithiothreitol (DTT), 1 mM PMSF, 1 mM benzamidine, 0.5 μg/mL LPP, 50 mM NaF, 10 mM β-glycerophosphate, 1 mM Na_3_VO_4_. Crude lysates were sonicated with a Hielscher handheld probe sonicator (5 sec., cycle 1, amplitude 70 %) and cleared at maximum speed via centrifugation (10 min., 21 000 x g, 4 °C). Lysates mixed with SDS sample buffer were loaded on a SDS PAGE gel and blotted as described above. Blots were analyzed for Rep and VP expression levels, as well as the phosphorylation state of KAP1^S824^.

gDNA and AAV2 viral genomes were extracted from the second cell pellet using the GenElute™ Mammalian Genomic DNA miniprep kit (Sigma-Aldrich). qPCR was performed to quantify the viral genomes per cell (vg/cell). A standard curve of HEK293T gDNA and cut pAV2 plasmid was included to allow the precise quantification of the number of cells and AAV2 genomes ^19^. For each sample, two different primer pairs for two different genes (cyclophilin and AAV2 *cap*) were used. All vg/cell values were normalized to the control group used in each experiment, either NT siRNA or pLKO empty vector. For primer pair sequences, see supplementary table.

### AlphaFold 3 multimer prediction

For AlphaFold 3 multimer predictions, the AlphaFold server from google DeepMind was used (https://alphafoldserver.com). Primary amino acid sequences of the proteins were obtained from Uniprot. Protein sequences were submitted on the AlphaFold server to model the interaction between the proteins. Models were analysed via UCSF ChimeraX.

## Acknowledgements

We thank Benjamien Moeyaert, Samir Nuseibeh, and Sofie De Munter for their valuable feedback on the written manuscript. This project received financial support from the Research Foundation Flanders (grant no. G0C3220N) and the KU Leuven Research Fund (3M190475). Bram Vandewinkel was supported by a PhD fellowship for strategic basic research from the Research Foundation Flanders (grant no. 1S58121N).

## Author contributions

B.V.: Conceptualization, supervision, experimental design, performance and analysis of the majority of experiments. B.V. also wrote the manuscript. S.T. generated the shRNA 3’-UTR PPP4C HEK293T cell line and obtained the data shown in figure 3C and 3F. Z.C. provided expert advice on the performance of the lysate-based split-luciferase assays and designed and validated the split-luciferase constructs in figure 5D. M.B.: conceptualization, supervision, funding acquisition and review. E.H.: conceptualization, supervision, funding acquisition, project administration, and writing—original draft, review, and editing. All authors read the final manuscript and agreed with its contents.

