## Supplementary figures for "The adeno-associated virus Rep proteins target PP4:SMEK1 by preventing substrate-recruitment"

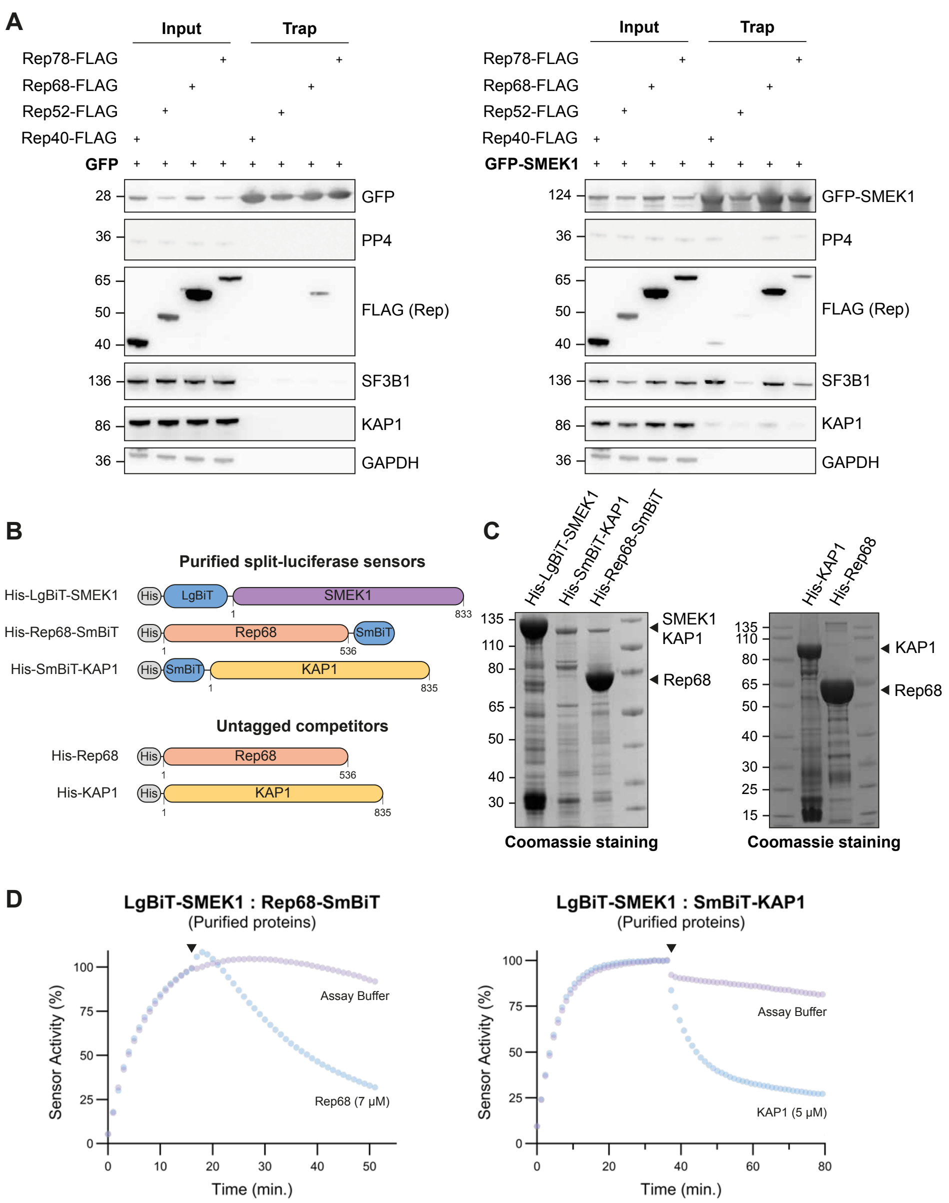


**Figure S1. Validation of the SMEK1:Rep interaction.**

**(A)** GFP-Trap experiment of GFP-tagged SMEK1 from cells ectopically expressing FLAG-tagged Rep (right panel). GFP expression alone served as a control (left panel). Input samples are shown on the left, while trap samples are shown on the right. **(B)** Design of the split-luciferase sensors and untagged competitors used for the protein-protein interaction studies with purified proteins. Sensors and competitors were tagged N-terminally with a 10X His-tag for recombinant expression and purification from *E. coli*. **(C)** Coomassie staining of the purified split-luciferase sensors and untagged competitors represented in **B** and used in **D**. **(D)** Kinetic-trace experiment with the

LgBiT-SMEK1:Rep68-SmBiT and LgBiT-SMEK1:SmBiT-KAP1 purified interaction sensors. 0.5 nM LgBiT-SMEK1 was mixed with 10 nM of SmBiT-tagged protein. The black arrow indicates the addition of untagged competitor. Concentration of the competitors is indicated in the graph. The presented data is plotted as a percentage of the signal-to-background (S/B) ratio right before the addition of competitor.

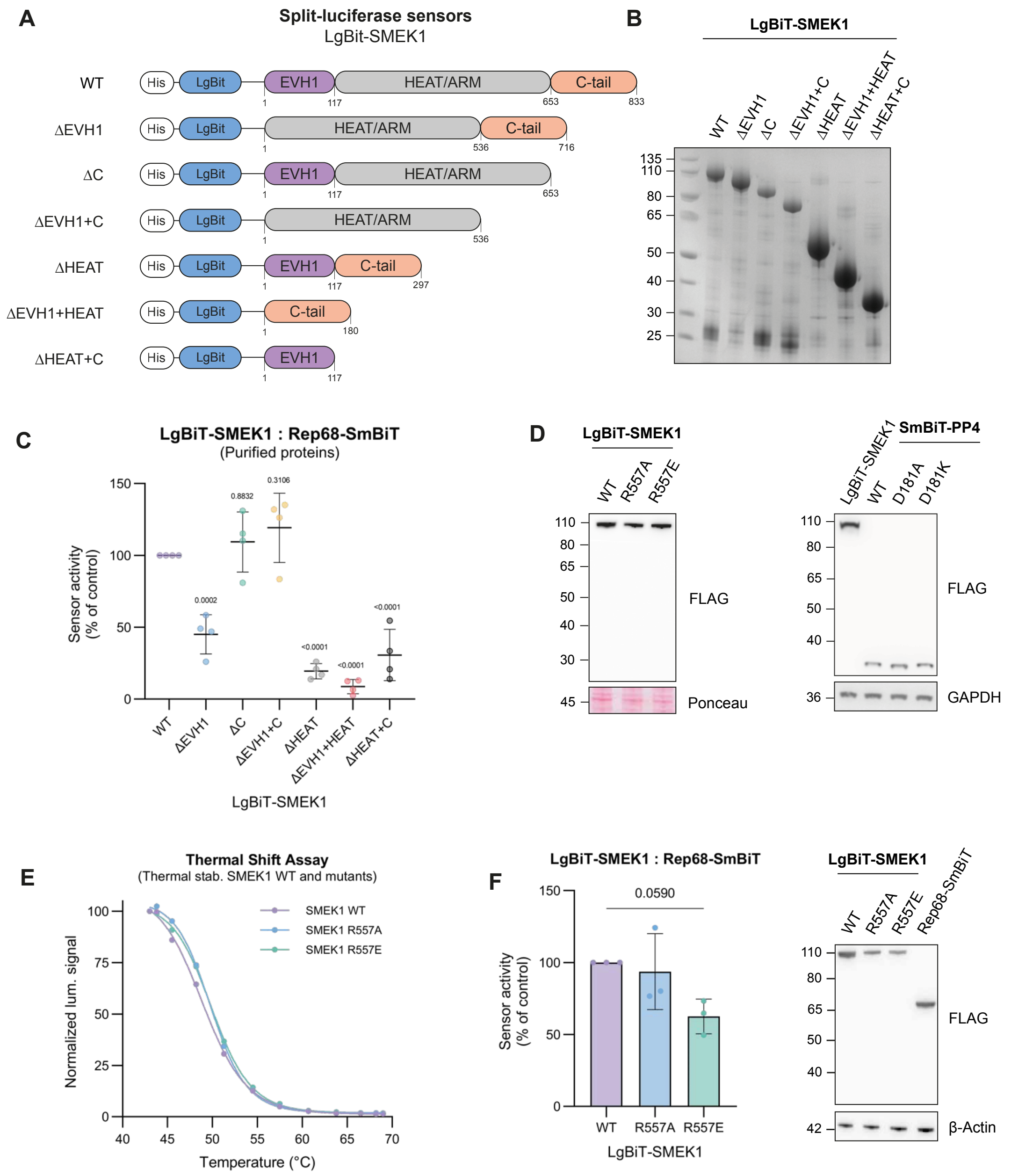


$T_m$:
- LgBit-SMEK1 WT = 48.98 °C
- LgBit-SMEK1 R557A = 49.84 °C
- LgBit-SMEK1 R557E = 49.95 °C

**Figure S2. The HEAT/Arm domain of SMEK1 is essential for Rep68 interaction.** **(A)** Design of the truncated SMEK1 split-luciferase sensors. Sensors were made for expression in HEK293T cells or *E. coli* (His-tagged). **(B)** Coomassie staining of the His-LgBiT-SMEK1 deletion mutants purified from *E. coli*. **(C)** Split-luciferase assays with the purified truncated LgBiT-SMEK1 (0.5 nM) and Rep68-SmBiT (10 nM) interaction sensors. End-point measurements were taken after mixing the sensor components and incubating them at room temperature for 20 minutes prior to read-out. **(D)** Immunoblot of the split-luciferase lysates used in figure 2F. **(E)** Thermal shift assay of WT and R557→A/E LgBiT-SMEK1 in lysates to assess the effect on the thermal stability of

SMEK1 upon introduction of a point mutation in the HEAT/Arm domain. **(F)** Lysate-based split-luciferase end-point measurement of the LgBiT-SMEK1$^{R557\rightarrow A/E}$:Rep68-SmBiT interaction sensors (left panel). Bioluminescence signal was measured after 20 minutes of incubation at room temperature. Data plotted as percentage of the LgBiT-SMEK1$^{WT}$:Rep68-SmBiT S/B signal. (mean ± SD; n = 3 independent repeats). Statistical significance was determined by a two-sided unpaired t-test. Immunoblot of the lysates is shown on the right.

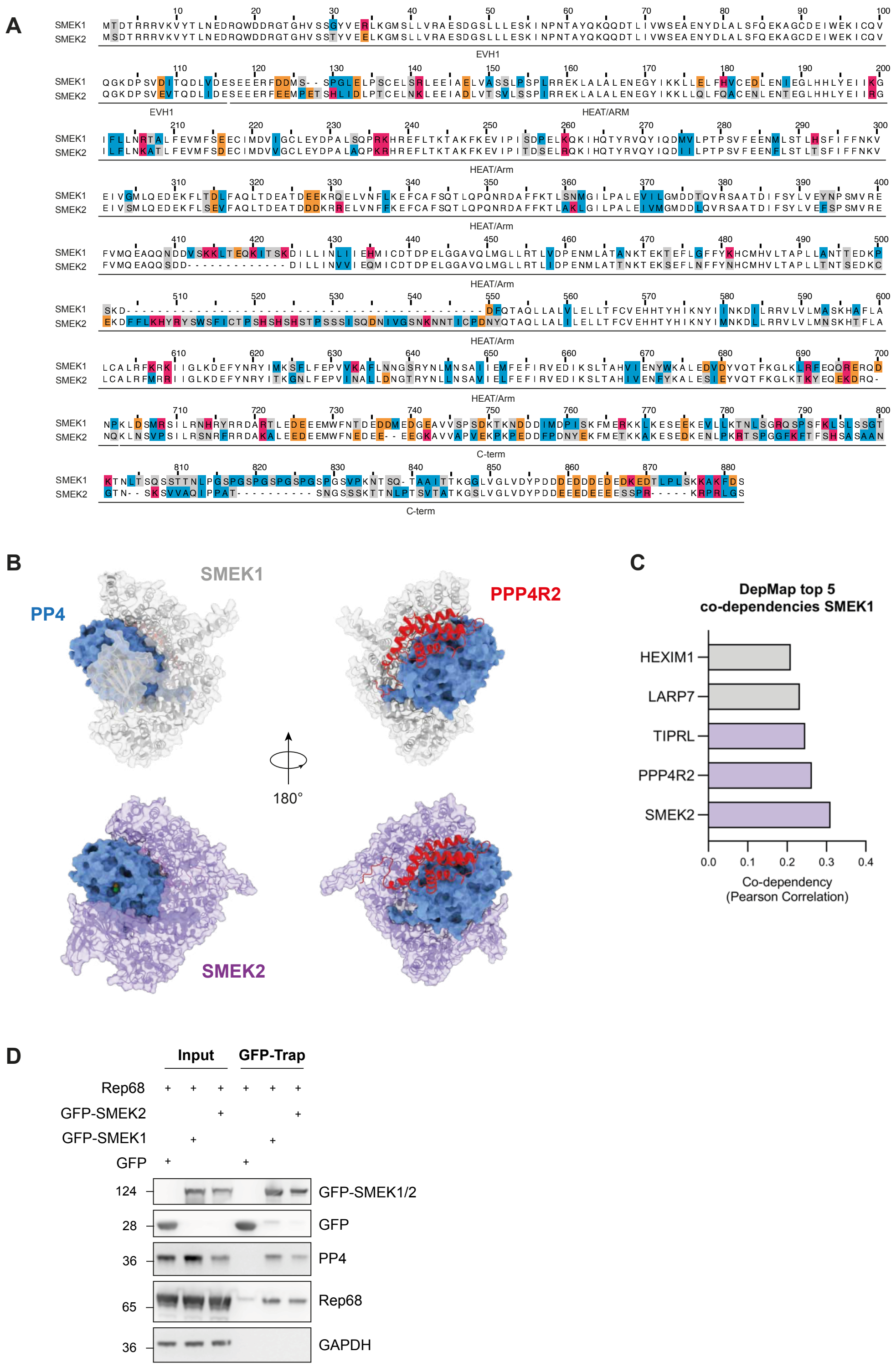


**Figure S3. SMEK1 and SMEK2 exhibit extensive sequence and structural homology.**

**(A)** Primary amino acid sequence alignment of SMEK1 and SMEK2, with only differing residues highlighted. Hydrophobic, negatively charged, positively charged, and other amino acids are color-coded in blue, orange, red, and grey, respectively. **(B)** AlphaFold 3 multimer predictions of the PP4:PP4R2:SMEK1 and PP4:PP4R2:SMEK2 heterotrimeric complexes, showing the structural homology between the two complexes. **(C)** DepMap CRISPR knockout data showing a functional dependency between SMEK1 and SMEK2. **(D)** GFP-trap of GFP-tagged SMEK1 and SMEK2 from cells co-expressing Rep68.

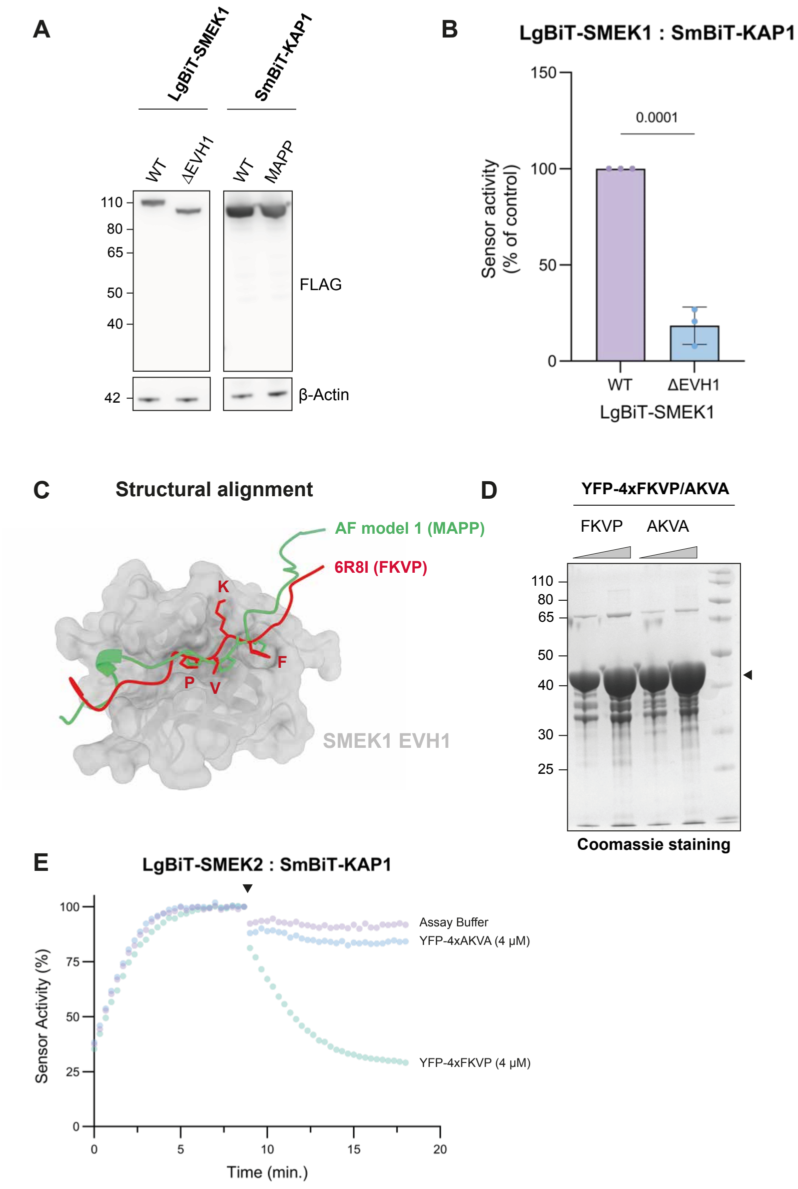


**Figure S4. The $^{423}$MAPP$^{426}$ SLiM motif of KAP1 interacts with SMEK1 and SMEK2.** **(A)** Immunoblot of the LgBiT-SMEK1$^{WT/\Delta EVH1}$ and SmBiT-KAP1$^{WT/MAPP}$ split-luciferase lysates. **(B)** Lysate-based split-luciferase end-point measurement of the LgBiT-SMEK1$^{WT}$:SmBiT-KAP1 and LgBiT-SMEK1$^{\Delta EVH1}$:SmBiT-KAP1 interaction sensors. Bioluminescence signal was read out after 25 minutes incubation at room temperature and plotted as a percentage of the LgBiT-SMEK1$^{WT}$:SmBiT-KAP1 signal (mean ± SD; n = 3 independent repeats). Statistical significance was determined by a two-sided unpaired t-test. **(C)** Structural alignment of the EVH1:FKVP co-crystal structure (PDB 6R8I) with the AlphaFold 3 model 1 prediction shown in figure 4C. **(D)** Coomassie staining of the purified YFP-4xFKVP and YFP-4xAKVA fusion proteins performed to assess the purity. **(E)** Kinetic-trace experiment of the LgBiT-SMEK2:SmBiT-KAP1 interaction sensor. Arrow indicates the addition of purified YFP-4xFKVP competitor or the AKVA control (concentrations indicated in the graph). The represented data is plotted as a percentage of the signal-to-background (S/B) ratio right before addition of the competitor.

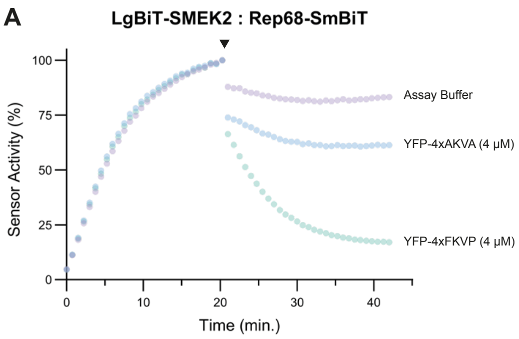


**Figure S5. Rep68 interacts with SMEK2 and is outcompeted with the FKVP peptide.**

Kinetic-trace experiment of the LgBiT-SMEK2:Rep68-SmBiT interaction sensor. Arrow indicates the addition of purified YFP-4xFKVP competitor or the AKVA control (concentrations indicated in the graph). The represented data is plotted as a percentage of the S/B ratio right before addition of the competitor.

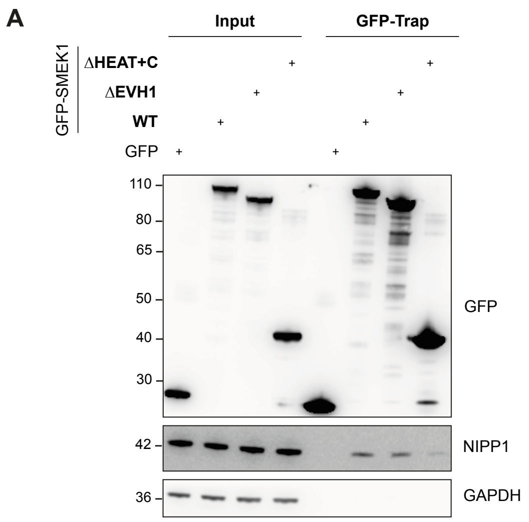


**Figure S6. The HEAT/arm and/or C-term of SMEK1 interacts with NIPP1.**

GFP-trap of GFP-tagged SMEK1 (WT, ΔEVH1, ΔHEAT+C-term) or GFP alone (control) to check for co-precipitation of endogenous NIPP1.
