## Supplementary materials for "The adeno-associated virus Rep proteins target PP4:SMEK1 by preventing substrate-recruitment"

| Recombinant DNA | | |
|---|---|---|
| **Plasmid** | **Source** | **Identifier** |
| Rep78-FLAG | This work | N/A |
| GFP-N1 | Addgene | Cat # 6085-1 |
| GFP-SMEK1 | This work | N/A |
| FLAG-LgBit-SMEK1 | This work | N/A |
| Rep68-SmBit-FLAG | This work | N/A |
| FLAG-SmBit-KAP1 | This work | N/A |
| FLAG-SMEK2 | This work | N/A |
| His-KAP1 | This work | N/A |
| His-Rep68 | This work | N/A |
| Rep68-FLAG | This work | N/A |
| Rep52-FLAG | This work | N/A |
| Rep40-FLAG | This work | N/A |
| His-FLAG-LgBit-SMEK1 | This work | N/A |
| His-Rep68-SmBit-FLAG | This work | N/A |
| His-FLAG-SmBit-KAP1 | This work | N/A |
| FLAG-LgBit-SMEK1 ΔEVH1 | This work | N/A |
| FLAG-LgBit-SMEK1 ΔC | This work | N/A |
| FLAG-LgBit-SMEK1 ΔEVH1+C | This work | N/A |
| FLAG-LgBit-SMEK1 ΔHEAT | This work | N/A |
| FLAG-LgBit-SMEK1 ΔEVH1+HEAT | This work | N/A |
| FLAG-LgBit-SMEK1 ΔHEAT+C | This work | N/A |
| His-FLAG-LgBit-SMEK1 ΔEVH1 | This work | N/A |
| His-FLAG-LgBit-SMEK1 ΔC | This work | N/A |
| His-FLAG-LgBit-SMEK1 ΔEVH1+C | This work | N/A |

| | | |
|---|---|---|
| His-FLAG-LgBit-SMEK1 ΔHEAT | This work | N/A |
| His-FLAG-LgBit-SMEK1 ΔEVH1+HEAT | This work | N/A |
| His-FLAG-LgBit-SMEK1 ΔHEAT+C | This work | N/A |
| FLAG-SmBiT-PP4 | This work | N/A |
| GFP-SMEK1 ΔEVH1 | This work | N/A |
| GFP-SMEK1 ΔHEAT+C | This work | N/A |
| FLAG-LgBit-SMEK1 R557A | This work | N/A |
| FLAG-LgBit-SMEK1 R557E | This work | N/A |
| FLAG-SmBiT-PP4 D181A | This work | N/A |
| FLAG-SmBiT-PP4 D181K | This work | N/A |
| FLAG-SMEK1 | This work | N/A |
| FLAG-SMEK1 R557A | This work | N/A |
| FLAG-SMEK1 R557E | This work | N/A |
| Rep68 | This work | N/A |
| Tet-pLKO-puro | Addgene | #21915 |
| pRSv REV | Addgene | #12253 |
| pMD2 VsVg | Addgene | #12259 |
| pMDL gag/pRRE #54 | Addgene | #12251 |
| MGC Human PPP4R3A Sequence-Verified cDNA | Horizon Discovery | MHS6278-202759611 |
| MGC Human PPP4C Sequence-Verified cDNA | Horizon Discovery | MHS6278-202827307 |
| MGC Human PPP4R3B Sequence-Verified cDNA | Horizon Discovery | MHS6278-202827304 |
| GFP-SMEK2 | This work | N/A |

| | | |
|---|---|---|
| FLAG-SmBit-KAP1 MAPP→AAAA | This work | N/A |
| His-YFP-4X(FxxP) | This work | N/A |
| His-YFP-4X(AxxA) | This work | N/A |
| FLAG-LgBiT-SMEK2 | This work | N/A |
| FLAG-LgBiT-B56 | Claes et al., 2023 | N/A |
| FLAG-SmBit-RepoMan | Claes et al., 2023 | N/A |
| YFP-4X(LSPI) | Claes et al., 2023 | N/A |
| FLAG-NIPP1 | Recieved from Prof. Mathieu Bollen | N/A |
| FLAG-NIPP1 Δ1-22 | This work | N/A |
| FLAG-NIPP1 ΔFHA | This work | N/A |
| GFP-SMEK1 TP→AA | This work | N/A |
| GFP-SMEK1 TP→DP | This work | N/A |

| Antibodies | | |
|---|---|---|
| **Details antibody** | **Source** | **Identifier** |
| Mouse monoclonal anti-FLAG® M2 | Sigma-Aldrich | Cat # F1804 |
| Mouse monoclonal anti-SAP155 | MBL Iinternational Corporation | Cat # D221-3 |
| Mouse monoclonal anti-TIF1b | Merck Millipore | Cat # MAB3662 |
| Rabbit polyclonal anti-SMEK1 | Abcam | Cat # ab70635 |
| Rabbit polyclonal anti-PPP1R8 | Atlas Antibodies | Cat # HPA027452 |
| Rabbit monoclonal anti-GAPDH | Cell Signalling technologies | Cat # 2118S |
| Mouse monoclonal anti-GFP | Santa Cruz | Cat # sc-9996 |
| Mouse monoclonal anti-AAV2 Rep | Progen | Cat # 65172 |
| Rabbit polyclonal anti-PPP4C | Invitrogen | Cat # PA596059 |
| Rabbit polyclonal anti-AAV VP1/2/3 | Progen | Cat # 61084 |
| Rabbit polyclonal anti-phospho-KAP1 (S824) | Bethyl Laboratories | Cat # A300-767A |
| Rabbit polyclonal anti-PPP4R3B | Atlas Antibodies | Cat # HPA001233 |
| Rabbit RPA2 | Abcam | Cat # ab76420 |
| Rabbit polyclonal anti-phospho-RPA2 (S4/8) | Invitrogen | Cat # PA5121338 |
| Rabbit polyclonal anti-phospho-RPA2 (S33) | Invitrogen | Cat # PA539809 |
| Mouse monoclonal anit-b-Actin | Sigma-Aldrich | Cat # A5441 |
| Mouse monoclonal anti-PP1 | Recieved from Prof. Mathieu Bollen | NA |

| siRNA | | |
|---|---|---|
| **Product** | **Source** | **Identifier** |
| ON-TARGETplus Human PPP4R3A siRNA SMARTPool | Horizon Discovery | L-019093-00-0020 |
| ON-TARGETplus Human PPP4R3B siRNA SMARTPool | Horizon Discovery | L-027284-00-0020 |
| ON-TARGETplus Non-targeting Control Pool | Horizon Discovery | D-001810-10-20 |

| Other products | | |
|---|---|---|
| **Product** | **Source** | **Identifier** |
| JetPRIME® | Polyplus | Cat # 101000046 |
| PEI MAX® | Polysciences | Cat # 24765-1 |
| NuPAGE 4-12 % Bis-Tris gels | Invitrogen | Cat # NW04125BOX |
| Q5 High-Fidelity Polymerase | NEB | Cat # M0491 |
| NEBuilder® HiFi DNA Assembly | NEB | Cat # E5520S |
| HisPur™ Ni-NTA Resin | Thermo Scientific™ | Cat # 88222 |
| ANTI-FLAG® M2 Affinity Gel | Millipore | Cat # A2220 |
| Anti-eGFP nanobody beads | Home made | N/A |

**Mammalian and bacterial cells**

| Product | Source | Identifier |
|---|---|---|
| HEK293T | Recieved from Prof. Mathieu Bollen | N/A |
| HEK293T shRNA PP4-3'UTR knockdown cell line | Home made | N/A |
| DH5alpha competent cells | Home made | N/A |
| BL21-Gold(DE3) competent cells | Home made | N/A |

**Software**

| Product | Source | Identifier |
|---|---|---|
| PyMOL | Schrödinger | https://pymol.org/ |
| UCSF ChimeraX | National Institute of Health | https://www.cgl.ucsf.edu/chimerax/ |
| ImageQuant LAS4000 | GE Healthcare | N/A |
| GraphPad Prism | GraphPad Software, LLC | https://www.graphpad.com/ |
| Image Lab 6.1 | Bio-Rad Laboratories Inc | https://www.bio-rad.com/ |
| Adobe Illustrator | Adobe | https://www.adobe.com/ |

**Chemicals**

| Product | Source | Identifier |
|---|---|---|

| | | |
|---|---|---|
| Western Lightning Plus-ECL, Enhanced Chemiluminescence Substrate | Perkin Elmer | Cat # NEL105001EA |
| Doxycycline | Sigma-Aldrich | Cat # 24390-14-5 |
| IPTG | | |

**Virus**

| **Product** | **Source** | **Identifier** |
|---|---|---|
| AAV serotype 2 | Home made | N/A |
| Adenovirus serotype 5 | Home made | N/A |

**shRNA**

| **Gene** | **Sequence (5'-3')** |
|---|---|
| PPP4C 3'-UTR | CCGGACCATGAAGTTTCCAATAATTCTCGAGAATTATTGGAAACTTCATGGTTTTTT<br>AATTAAAAAACCATGAAGTTTCCAATAATTCTCGAGAATTATTGGAAACTTCATGGT |

**qPCR primers**

| **Gene** | **Sequence (5'-3')** |
|---|---|
| Cyclophilin | GGACGGACATTTTCACCCCT<br>GTCCCGTGGAGTACTGTGTG |
| AAV cap | TGCTGGACCCAACACAAATG<br>TGCCATCCAACCACTCAGTCT |
